# A stress-induced Tyrosine tRNA depletion response mediates codon-based translational repression and growth suppression

**DOI:** 10.1101/416727

**Authors:** Doowon Huh, Maria C. Passarelli, Jenny Gao, Shahnoza N Dusmatova, Clara Goin, Lisa Fish, Alexandra M. Pinzaru, Henrik Molina, Elizabeth A. McMillan, Hosseinali Asgharian, Hani Goodarzi, Sohail F. Tavazoie

## Abstract

Eukaryotic transfer RNAs can become selectively fragmented upon various stresses, generating tRNA-derived small RNA fragments. Such fragmentation has been reported to impact a small fraction of the tRNA pool and thus presumed to not directly impact translation. We report that oxidative stress can rapidly generate tyrosine tRNA_GUA_ fragments in human cells—causing significant depletion of the precursor tRNA. Tyrosine tRNA_GUA_ depletion impaired translation of growth and metabolic genes enriched in cognate tyrosine codons. Depletion of tyrosine tRNA_GUA_ or its translationally regulated targets USP3 and SCD repressed proliferation—revealing a dedicated tRNA-regulated growth suppressive pathway for oxidative stress response. Tyrosine fragments are generated in a DIS3L2 exoribonuclease-dependent manner and inhibit hnRNPA1-mediated transcript destabilization. Moreover, tyrosine fragmentation is conserved in *C. elegans*. Thus, tRNA fragmentation can coordinately generate *trans*-acting small-RNAs and functionally deplete a tRNA. Our findings reveal the existence of an underlying adaptive codon-based regulatory response inherent to the genetic code.

## INTRODUCTION

Transfer RNAs (tRNAs) are universal decoders of the genetic code. By recognizing three-nucleotide sequences (codons) in transcripts, tRNAs enable ribosomal incorporation of specific amino acids into the growing polypeptide chain. Because there exists a larger number of codons than amino acids, the code is degenerate, with multiple ‘synonymous’ codons encoding a given amino acid. The human genome contains over 400 tRNA genomic loci, with multiple genes encoding tRNAs that contain the same anti-codon (Parisien et al., 2013). Such tRNA molecules that recognize the same trinucleotide codon sequence are termed tRNA isodecoders. The central roles of tRNAs in translation have been defined through elegant structural and biophysical studies (Nissen et al., 2000; Ogle et al., 2002). Recent studies have, however, challenged our textbook notions regarding these essential molecules and suggest that beyond their static roles as adaptors in translation, tRNAs play additional dynamic roles in gene regulation (Schimmel, 2018).

One line of support for non-canonical roles for tRNAs was the discovery in *Tetrahymena* that starvation stress caused a subset of tRNAs to undergo endonucleolytic cleavage into smaller fragments (Lee and Collins, 2005). In humans, tRNA-derived fragments (tRFs) were originally detected in the urine of cancer patients four decades ago but were of unknown function (Gehrke et al., 1979). A variety of stresses have since been observed to generate tRFs in cells of organisms ranging from yeast to man (Thompson et al., 2008). Specific ribonucleases have also been implicated in generating tRFs in distinct species, including Rny1p in yeast and angiogenin (ANG) in human cells (Fu et al., 2009; Thompson and Parker, 2009a). Past studies reported that tRNA fragmentation did not noticeably deplete (<1%) the precursor tRNA molecules of the specific tRFs being studied (Saikia et al., 2012). Such observations suggested that the primary consequence of tRF generation was *trans* action by these small RNAs rather than translational impairment owing to depletion of the precursor tRNA. Consistent with this hypothesis, endogenous tRFs have been found to interact in *trans* with RNA-binding proteins and to mediate post-transcriptional gene repression (Goodarzi et al., 2015; Kuscu et al., 2018). Attesting to their functional roles, inhibition of tRFs has been shown to impact malignant phenotypes at the cellular and organismal levels, while deletion of ANG elicited defects in hematopoiesis (Goncalves et al., 2016).

A second line of evidence for non-canonical roles by tRNAs was the observation that the expression levels of some tRNAs become modulated in the context of malignancy and cancer progression (Goodarzi et al., 2016; Pavon-Eternod et al., 2009). Such observations challenged the common dogma that tRNAs are static components in mammalian cells. The advent of tRNA microarray methods enabled assessing the levels of large numbers of tRNAs in normal and malignant cells (Dittmar et al., 2006). Comparison of breast cancer cells to non-malignant breast cells revealed a number of tRNAs to be overexpressed and others repressed, perhaps owing to genomic instability and subsequent copy number alterations of tRNA genes (Goodarzi et al., 2016; Pavon-Eternod et al., 2009). Such alterations in tRNA content have been associated with altered cellular protein expression (Chan et al., 2010; Pavon-Eternod et al., 2013; Pershing et al., 2015) and mRNA stability (Boel et al., 2016; Hoekema et al., 1987; Presnyak et al., 2015). Other analyses revealed similar findings of large-scale tRNA expression alterations associated with the cancerous proliferative state versus the differentiated state (Gingold et al., 2014). Studies of cancer progression revealed that upregulation of specific tRNA isodecoders through genomic copy number gains causally enhanced translation of specific proteins that promoted cellular invasiveness and metastatic capacity (Goodarzi et al., 2016). Mutagenesis studies of the KRAS oncogene also provided support for distinctness among ‘synonymous’ codons; KRAS protein expression became upregulated upon mutation of a ‘rare’ codon (decoded by a low-abundant tRNA isoacceptor) to an ‘optimal’ synonymous codon (decoded by an abundant tRNA isoacceptor) (Pershing et al., 2015)—consistent with tRNA availability impacting protein expression (Gustafsson et al., 2004). If distinct isodecoders can modulate different sets of genes and elicit distinct phenotypes, then one may expect that a mutation in a tRNA isodecoder could give rise to a specific phenotype. Indeed, genetic studies revealed that a mutation in the anticodon of a single nuclear encoded mouse tRNA gene could cause a specific phenotype—cerebellar neurodegeneration (Ishimura et al., 2014).

The observed disease-associated tRNA modulations and the ensuing codon-dependent gene expression effects in the context of genomic instability and cancer raise the question of whether such tRNA modulations occur in normal cells and perhaps elicit translational effects in response to exogenous signals. We herein describe the existence of an endogenous gene regulatory response to oxidative stress in human cells that is mediated by the depletion of a specific tRNA. We observed that oxidative stress rapidly induced fragmentation of tyrosine tRNA_GUA_—leading to tRNA^Tyr^_GUA_ depletion and reduced expression of a gene set enriched in tyrosine codons. The affected genes were significantly enriched in growth and metabolic pathways. The depletion of tRNA^Tyr^_GUA_ and the resulting impaired translation of downstream genes caused growth suppression. Moreover, we observe that the tyrosine tRF^Tyr^_GUA_ that is produced interacts in *trans* with the hnRNPA1 and SSB RNA binding proteins. Oxidative-stress induced RNA^Tyr^_GUA_ fragmentation is dependent on DIS3L2, an exoribonuclease associated with Perlman congenital overgrowth syndrome and Wilms tumor. Oxidative stress-induced tyrosine tRNA_GUA_ fragmentation is also conserved in the nematode *C. elegans*. Our findings uncover a direct relationship between tRNA fragmentation and tRNA modulation and reveal that for a specific tRNA, stress-induced fragmentation can be substantial enough to deplete the tRNA precursor, with adaptive translational gene regulatory consequences.

## RESULTS

### Systematic characterization of changes in tRNAs and tRFs as a response to oxidative stress

We hypothesized that stress-induced tRNA fragmentation may deplete the pools of specific tRNAs. To search for such tRNAs, we sought to globally profile changes in tRNA and tRF levels upon exposure to oxidative stress—a stress known to robustly induce tRF generation (Thompson et al., 2008; Thompson and Parker, 2009a; Yamasaki et al., 2009). Due to numerous nucleotide modifications as well as their highly conserved secondary structures, tRNAs are poor substrates for reverse transcription and consequently are poorly quantified by standard high-throughput sequencing methods. To overcome this, multiple independent methods have been developed to profile tRNA species (Cozen et al., 2015; Gogakos et al., 2017; Goodarzi et al., 2016; Zheng et al., 2015). We employed one such method—a probe-based tRNA capture, ligation, and deep-sequencing approach—to profile tRNA isodecoder abundances across samples (Goodarzi et al., 2016). This allowed us to assess the impact of oxidative stress on tRNA levels in a human mammary epithelial cell line (MCF10A). At both 8 and 24 hours after 200uM hydrogen peroxide (H_2_O_2_) treatment, tRNA levels were globally similar to tRNA levels in the non-treated condition (Fig 1A). However, multiple specific tRNAs (tRNA^Tyr^_GUA_, tRNA^Ile^_UAU_, tRNA^Leu^_UAA_, tRNA^Thr^_AGU_, and others) became significantly decreased in abundance over time after H_2_O_2_ treatment. We next exposed MCF10A cells to oxidative stress and profiled small RNA (smRNA) abundances via deep-sequencing (Fig 1B). This revealed oxidative-stress induction of many tRFs—consistent with previous studies describing the effects of oxidative stress on fragmentation of specific tRNAs by northern blot analyses (Saikia et al., 2012; Thompson et al., 2008). The observed tRF inductions were not artifacts of cell death, as cell viability was unchanged in treated versus control samples at the concentration of H_2_O_2_ used (Fig S1A). Integration of tRF sequencing and tRNA profiling analyses identified a set of candidate tRNAs that became fragmented and depleted upon oxidative stress (Fig 1C). From this overlapping set, tRNA^Tyr^_GUA_, tRNA^Leu^_UAA_, tRNA^Leu^_CAG_ exhibited significantly higher levels of oxidative stress-induced tRF generation (Fig 1B, Fig S1B).

**Figure 1.**
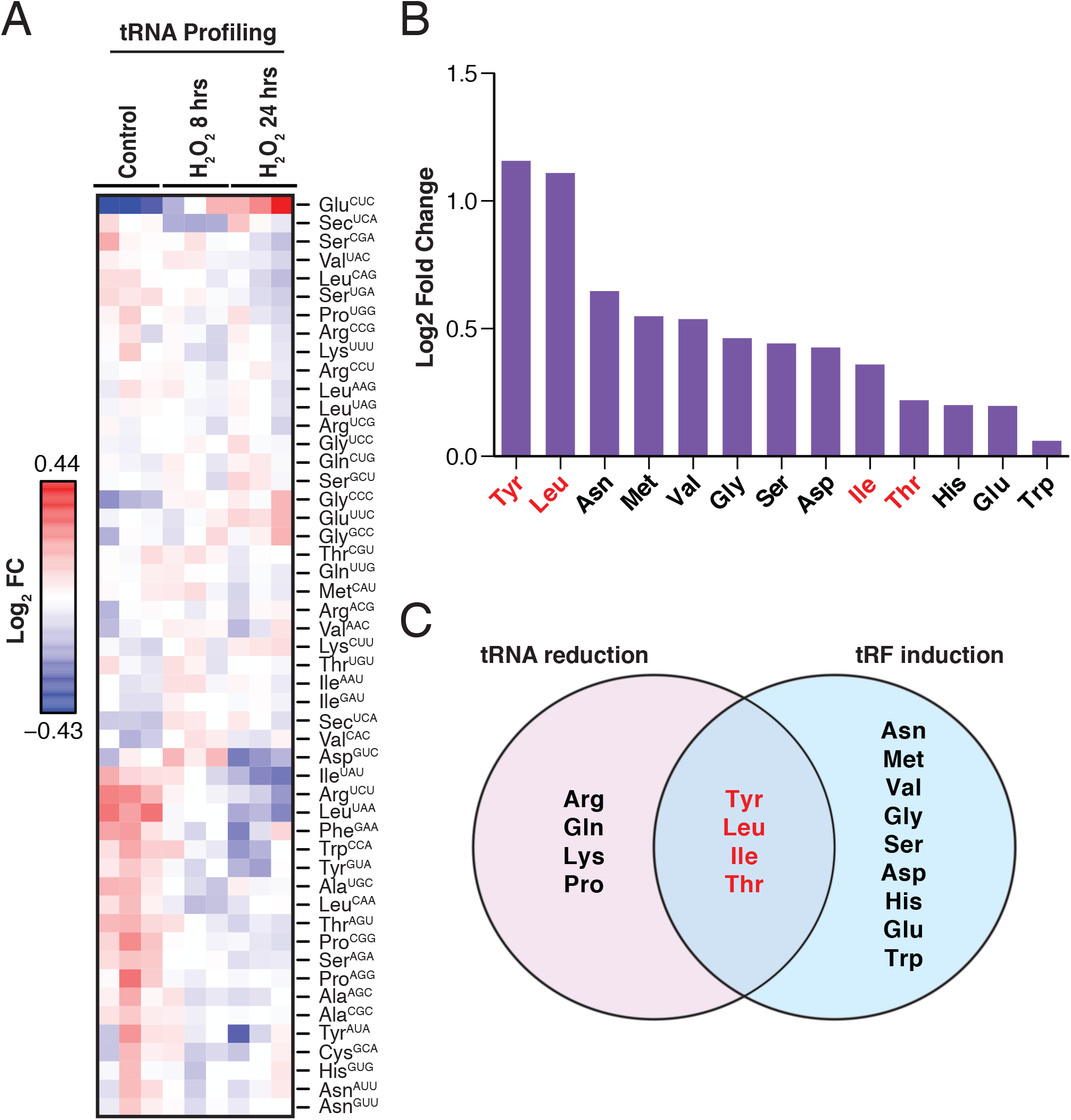
Identification of modulated mammalian tRNAs and tRFs in response to oxidative stress. (A) Heatmap of tRNA profiling of MCF10A cells at 8 and 24 hours post exposure to oxidative stress (200μM H_2_O_2_). Biological triplicate data is depicted at each time point relative to control cells. (B) MCF10A cells were exposed to oxidative stress (200μM H_2_O_2_) and processed for small RNA-sequencing. The log_2_-fold induction levels for tRFs derived from distinct tRNA isoacceptor is plotted. (C) Schematic depicts the overlap of tRNAs in (A) that decreased over time with tRFs from (B) that were induced. Four isoacceptor families of tRNAs are shown in the overlap with tyrosine-tRNA and leucyl-tRNAs as the most promising candidates that exhibited the highest degree of tRF induction.

### Oxidative stress-induced fragmentation depletes tRNA^Tyr^_GUA_

Northern blot analysis confirmed that oxidative stress can induce generation of tRF^Tyr^_GUA_ and tRF^Leu^ (multiple isodecoders). We thus focused our efforts on the tRF that exhibited the strongest induction by northern blot (tRF^Tyr^_GUA_) and its associated tRNA (Fig S2A). To determine if tRF^Tyr^_GUA_ formation is conserved in the nematode, we exposed *C. elegans* to H_2_O_2_. Similar to our observations in human cells, we observed generation of tRF^Tyr^_GUA_ following brief exposure (15 minutes) of *C. elegans* to H_2_O_2_ (Fig S2B)—suggesting conservation of this tRNA fragmentation response. To better characterize this response, we performed time-course studies in human cells. TRF^Tyr^_GUA_ became rapidly induced (within 5 minutes) upon cellular exposure to H_2_O_2_ (Fig 2A). TRF^Tyr^_GUA_ generation was associated with a concomitant precipitous decline in pre-tRNA^Tyr^_GUA_ levels, which remarkably became nearly undetectable at 1-hour post treatment (Fig 2B). This suggests that the majority of tRF^Tyr^_GUA_ is generated from the pre-tRNA^Tyr^_GUA_ rather than the mature tRNA, similar to our observations in *C. elegans*. Pre-tRNA-derived tRFs have been previously detected in other contexts (Lee et al., 2009). As tRNAs are one of the most stable classes of RNAs, with relatively long half-lives, we would expect a delayed effect on the abundance of the mature tRNA pool upon acute reduction of the pre-tRNA pool. Indeed, we observed a significant delayed reduction in the tRNA^Tyr^_GUA_ pool, which was observed at 24 hours post H_2_O_2_ exposure (Fig 2A-B). The mature tRNA^Tyr^_GUA_ pool diminished to roughly half the pre-treatment levels at 24 hours post treatment. These observations reveal that oxidative stress-induced generation of tRF^Tyr^_GUA_ can significantly deplete the corresponding mature tRNA^Tyr^_GUA_ pool. Our findings furthermore demonstrate that a single cellular stress can induce the levels of a specific tRF and deplete its corresponding tRNA.

**Figure 2.**
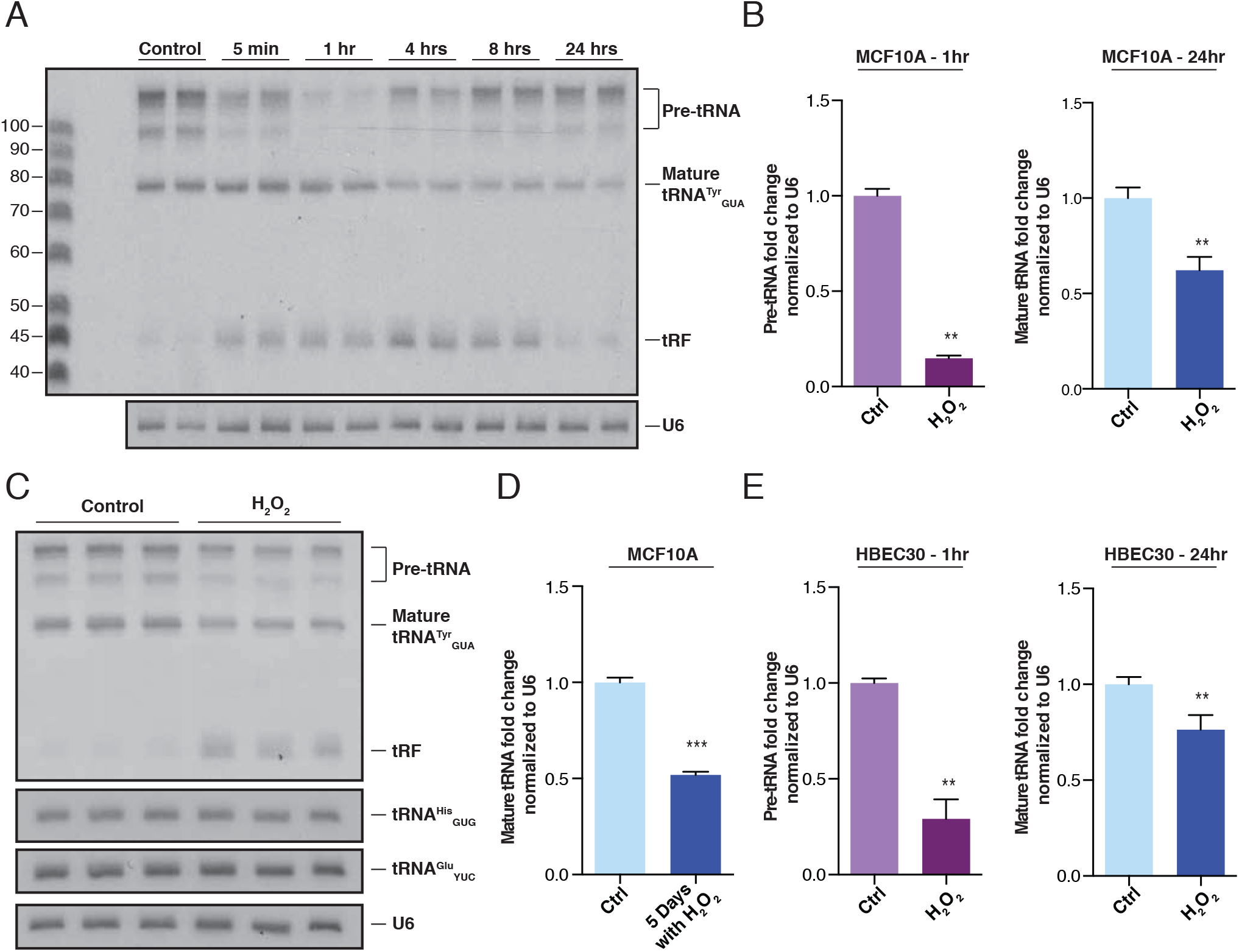
TRNA^Tyr^_GUA_ abundance is reduced while the corresponding Tyr-tRF is induced in response to oxidative stress. (A) A northern blot depicting a time course experiment ranging from five minutes to 24 hours of MCF10A cells in response to oxidative stress. A single probe complementary to pre-tRNA^Tyr^_GUA_, mature tRNA^Tyr^_GUA_, and tRF^Tyr^_GUA_ expression was ^32^P-labeled and used for detection. (B) Quantification of pre-tRNA^Tyr^_GUA_ northern blot analysis from multiple independent experiments after one hour as well as mature tRNA^Tyr^_GUA_ levels after 24 hours (normalized to U6 levels) are shown (n=6). (C) MCF10A cells were exposed to oxidative stress (200μM H_2_O_2_) once daily for five continuous days to test if repeated exposure to the stress could elicit a response similar to that found in (A). (D) Quantification of mature tRNA^Tyr^_GUA_ bands by northern blot after cells were treated once daily for five continuous days (normalized to U6) from multiple independent experiments (n=12). (E) Quantification of northern blot analysis for pre-tRNA^Tyr^_GUA_ (left) and tRNA^Tyr^_GUA_ (right) after one hour and 24 hours respectively in HBEC30 cells upon exposure to oxidative stress (200μM H_2_O_2_) as in (A) (n=6). Data represent mean ± s.e.m. A one-tailed Mann-Whitney test (*p < 0.05 and **p < 0.01) was used to test for statistical significance between the treated and control cell lines for each time point.

We next determined whether oxidative stress-induced tRNA^Tyr^_GUA_ depletion is a transient response or whether it could persist upon continual exposure to stress. Treatment of cells with H_2_O_2_ once daily for five consecutive days maintained tRNA^Tyr^_GUA_ repression as assessed by northern blot analysis (Fig 2C-D). RNA was collected on the sixth day, 24 hours after the last H_2_O_2_ exposure, and as before, tRF^Tyr^_GUA_ induction was maintained over this time (Fig 2C-D). Such continuous H_2_O_2_ treatment repressed mature tRNA^Tyr^_GUA_ levels by roughly half. We also assessed the effect of oxidative stress on a distinct mammalian cell line, the human bronchial epithelial cell line (HBEC30), and observed that pre-tRNA^Tyr^_GUA_ levels dramatically declined at 1-hour post treatment, with an ensuing significant reduction in mature tRNA^Tyr^_GUA_ levels at 24 hours (Fig 2E, Fig S2C). Despite this reduction in mature tRNA^Tyr^_GUA_ levels following H_2_O_2_ exposure, other control tRNAs—tRNA^His^_GUG_ and tRNA^Glu^_YUC_—remained unchanged relative to the control (Fig 2C, Fig S2D-E). To further confirm that tRNA^Tyr^_GUA_ levels are dependent on exposure to oxidative stress, we used an independent source of oxidative stress. Menadione, a commonly used pharmacological agent that induces oxidative stress, yielded similar results to that of H_2_O_2_. A reduction of the pre-tRNA and induction of the tRF were noted at early time points and mature tRNA^Tyr^_GUA_ levels significantly declined by 24 hours (Fig S2F-G). These findings reveal that stress-induced tRNA depletion can be a sustained response and that it can be elicited in multiple human cell types.

### Oxidative stress-induced tRNA^Tyr^_GUA_ depletion represses cellular growth

A possible explanation for the observed tRNA depletion effect could be that it is caused by cell death. However, H_2_O_2_ at the dose used in this study did not reduce cell viability (Fig S1A). Moreover, we observed depletion of specific tRNAs rather than a global effect on all tRNAs, which would be expected if this was a consequence of a non-specific cell death phenomenon. H_2_O_2_ doses at intermediate and sublethal ranges have been shown to cause growth arrest without cell death (Martindale and Holbrook, 2002). Consistent with such prior observations, we also observed impaired growth upon H_2_O_2_ treatment (Fig 3A). We thus sought to determine if tRNA^Tyr^_GUA_ depletion might contribute to growth repression as a response to oxidative stress. To determine the impact of reduced tRNA^Tyr^_GUA_ activity on cell growth, we undertook two orthogonal approaches. The first was to deplete tRNA^Tyr^_GUA_ through short-hairpin RNA-induced inactivation (Goodarzi et al., 2016). Using this approach, we generated a stable MCF10A cell line in which endogenous tRNA^Tyr^_GUA_ was depleted to roughly the same level as that observed upon H_2_O_2_ treatment of cells (Fig 3B). As an independent loss-of-function approach, we sought to impair cellular utilization of tRNA^Tyr^_GUA_ via RNAi-mediated depletion of its cognate amino acid charging enzyme—the tyrosine-tRNA synthetase (YARS) gene (Fig 3C). Due to guanine-uracil wobble base pairing (Crick, 1966; Ladner et al., 1975; Quigley and Rich, 1976), tRNA^Tyr^_GUA_ can recognize both codons (UAC and UAU) that code for the amino acid tyrosine. Moreover, we could not detect the other tyrosine tRNA (tRNA^Tyr^_AUA_) in MCF10A cells by northern blot, suggesting that this synonymous tRNA may play a minimal, if any, role in translation in these cells and is a rare tRNA—as has been reported by others in additional mammalian cell types (dos Reis et al., 2004). We reasoned that depletion of aminoacylated tRNA^Tyr^_GUA_ caused by YARS knockdown would phenocopy tRNA^Tyr^_GUA_ depletion since in both cases tRNA^Tyr^_GUA_–mediated tyrosine incorporation into proteins is impaired. We observed that impairment of tRNA^Tyr^_GUA_ function by either tRNA^Tyr^_GUA_ depletion or YARS depletion using two independent hairpins strongly impaired growth of MCF10A cells (Fig 3D). These results reveal that depletion of tRNA^Tyr^_GUA_ or inhibition of its cognate charging enzyme can significantly impair growth and phenocopy the H_2_O_2_ -induced physiological depletion of endogenous tRNA^Tyr^_GUA_. In contrast, overexpression of tRNA^Tyr^_GUA_ led to the opposite phenotype—increased cell growth (Fig 3E, Fig S3B). We propose that stress-induced depletion of tRNA^Tyr^_GUA_ constitutes an endogenous growth-suppressive stress response that contributes in part to the cellular oxidative stress response.

**Figure 3.**
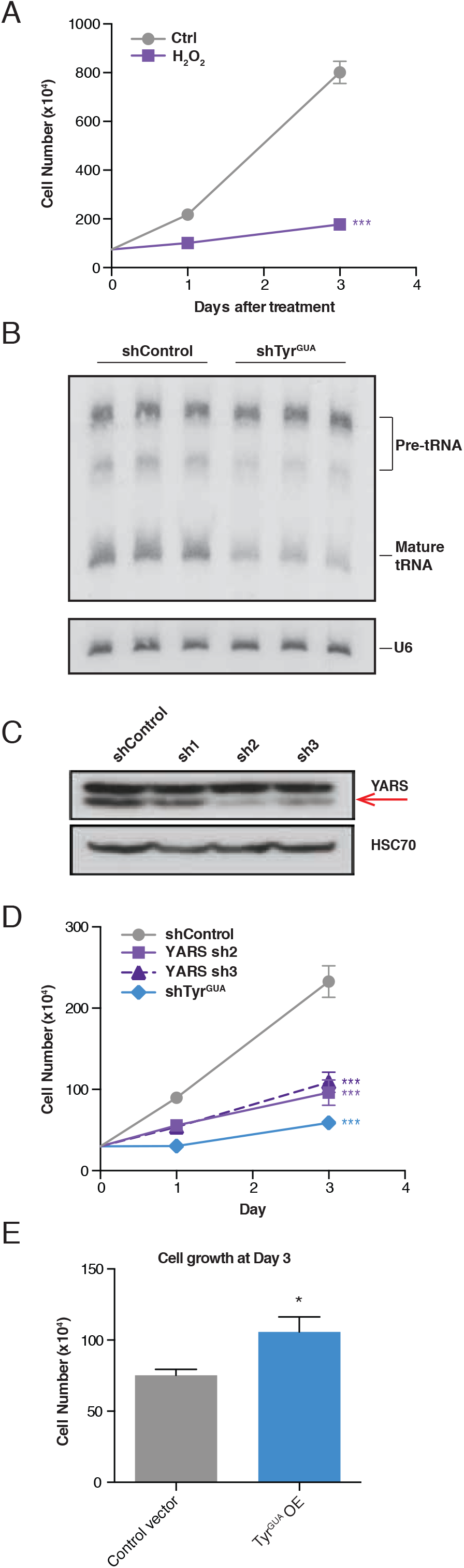
Cell growth repression upon oxidative stress and tRNA^Tyr^_GUA_ depletion. (A) Growth curves of MCF10A cells exposed to oxidative stress (200μM H_2_O_2_) relative to control cells (n=3). Two-way ANOVA was used to test for significance. (B) Northern blot of MCF10A cells expressing a control short-hairpin RNA or a hairpin targeting tRNA^Tyr^_GUA_. (C) A western blot of MCF10A expressing a control short-hairpin RNA or a hairpin targeting the tyrosine-tRNA synthetase, YARS (red arrow). HSC70 was used as a loading control. (D) Growth curves of MCF10A cells expressing RNAi against mature tRNA^Tyr^_GUA_ or YARS relative to cells expressing a control hairpin (n=3). Two-way ANOVA was used to test for significance. (E) Cell growth of MCF10A cells transiently transfected with a tRNA^Tyr^_GUA_ overexpression vector relative to an empty control vector (n=3). A one-tailed Mann-Whitney test was used to test for significance at day 3. Data represent mean ± s.e.m. *p < 0.05, **p < 0.01, and ***p < 0.001

### TRNA^Tyr^_GUA_ depletion represses expression of a set of growth genes

We hypothesized that stress-induced tRNA^Tyr^_GUA_ depletion impairs growth by reducing production of proteins enriched in its corresponding tyrosine codons. To search for such proteins, we conducted quantitative proteomic profiling of cells depleted of tRNA^Tyr^_GUA_ or impaired in tRNA^Tyr^_GUA_ aminoacylation. Label free mass-spectrometric proteomic profiling of cells depleted of tRNA^Tyr^_GUA_ or of YARS via shRNA-mediated knockdown revealed a highly significant correlation (R=0.648; p < 2.2e-16) in the proteomic profiles of these depleted cells relative to control hairpin expressing cells—consistent with a common set of downstream genes being impacted by these orthogonal methods of tRNA^Tyr^_GUA_ loss-of-function (Fig S4A). Amongst proteins that became depleted upon both of these perturbations, we searched for those that were also enriched for Tyr codons and are therefore likely to be tRNA^Tyr^_GUA_-dependent (Fig 4A). Using this approach, we identified 109 tyrosine-enriched proteins that exhibited sensitivity to tRNA^Tyr^_GUA_ depletion. This set of proteins was most significantly enriched in gene ontology (GO) functional categories (Ashburner et al., 2000; The Gene Ontology, 2017) associated with cellular growth, including regulation of ATP synthesis, G0 to G1 cell-cycle progression, and phosphorylation (Fig 4B). These findings reveal that tRNA^Tyr^_GUA_ depletion represses the abundance of a set of proteins associated with growth.

**Figure 4.**
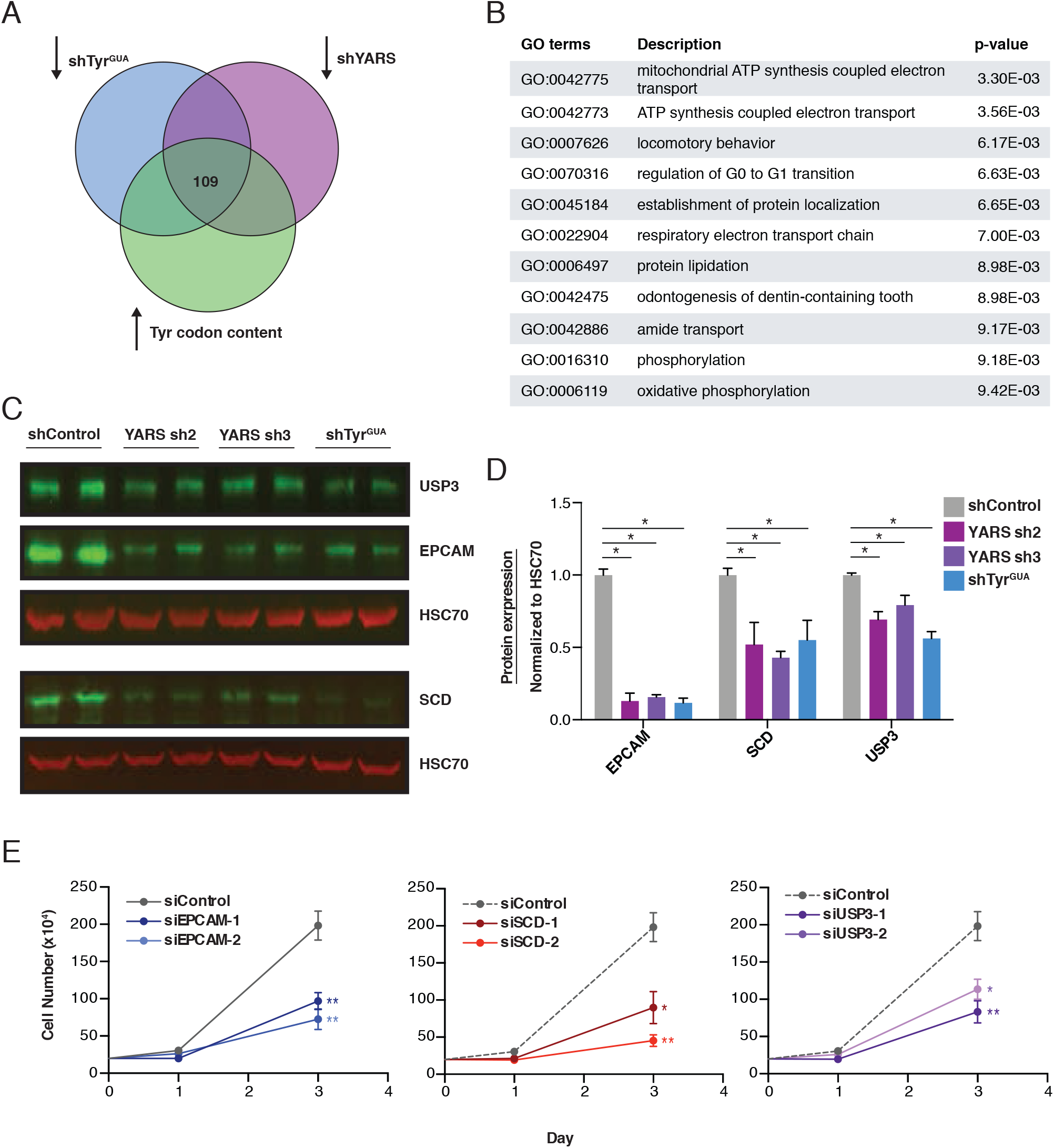
A set of growth promoting genes are sensitive to tRNA^Tyr^_GUA_ depletion. (A) Cells depleted of tRNA^Tyr^_GUA_ or YARS were processed for label free quantitation by mass spectrometry to identify proteins that were reduced by a log_2_-fold change of 0.5 or more. This set was overlapped with proteins containing a higher than median abundance of Tyr codon content to identify candidate mediators of the pleiotropic effects of tRNA^Tyr^_GUA_ depletion. (B) GO functional analysis of the 109 candidate gene-set from (A). (C) Quantitative western blot validation depicting abundances of protein targets (EPCAM, SCD, and USP3) identified from (A). HSC70 was used as a loading control and is not modulated upon molecular perturbation of tRNA^Tyr^_GUA_. (D) Quantification of western blot analysis in (C) (n=4). A one-tailed Mann-Whitney test was used to test for statistical significance between knockdown and control conditions. (E) Growth curves for MCF10A cells were transfected with either control siRNA or two independent siRNA targeting EPCAM, SCD, or USP3. Note that the control cell growth curve is the same in all graphs and were plotted separately for clarity and does not represent independent experiments. Two-way ANOVA was used to test for significance. Data represent mean ± s.e.m. *p < 0.05, **p < 0.01, and ***p < 0.001

We selected a small set of tRNA^Tyr^_GUA_–regulated genes that exhibited some of the greatest fold reductions upon tRNA^Tyr^_GUA_ depletion for further functional studies (Fig S4A-B). These genes comprised ubiquitin specific protease 3 (USP3), a hydrolase that deubiquitinates histone H2A and H2B and is required for mitotic entry and S phase progression (Nicassio et al., 2007), epithelial cell adhesion molecule (EPCAM), a tumorigenic cell surface protein over-expressed in many carcinomas (Munz et al., 2009), and stearoyl-CoA desaturase (SCD), the rate limiting enzyme for generating mono-unsaturated fatty acids such as palmitoleate and oleate—principle components of membrane phospholipids and cholesterol esters (Paton and Ntambi, 2009). Quantitative western blotting and quantitative real time PCR (qRT-PCR) revealed significant reductions in the endogenous levels of these proteins and mRNA transcripts upon shRNA-mediated depletion of tRNA^Tyr^_GUA_ or YARS (Fig 4C-D, Fig S4C). In contrast, the control protein HSC70 was not significantly depleted, consistent with our proteomic findings of a specific set of proteins being modulated upon tRNA^Tyr^_GUA_ depletion. To determine if repressed expression of these genes indeed impairs proliferation, we depleted these genes via RNA-interference. Knockdown of each of these genes by two independent hairpins repressed the growth of MCF10A cells—consistent with growth-promoting roles (Fig 4E, Fig S4D). Our results reveal that repressing the function of a specific tRNA by depleting it or inhibiting its aminoacylation, and thus its use in translation, represses expression of a set of tyrosine enriched proteins enriched in growth-dependent processes. Moreover, depletion of tRNA^Tyr^_GUA_ or its downstream regulated genes impairs growth of breast epithelial cells. We propose that this network constitutes a pro-growth tRNA^Tyr^_GUA_-regulated pathway and its repression via oxidative stress-induced tRNA fragmentation and depletion constitutes an adaptive growth suppressive stress response.

### TRNA^Tyr^_GUA_ depletion impairs protein translation in a codon-dependent manner

Our findings indicate that cellular tRNA^Tyr^_GUA_ expression regulates the abundance of a set of growth-associated proteins enriched in tyrosine codons. We hypothesized that these proteins would also be sensitive to oxidative stress-induced tRNA^Tyr^_GUA_ depletion. To test if these tRNA^Tyr^_GUA_-regulated proteins become repressed upon oxidative stress, we performed quantitative western blotting 24 hours after H_2_O_2_ treatment, a time point when tRNA^Tyr^_GUA_ is depleted. Consistent with our previous experiments, we noted significant reductions in protein levels of these tRNA^Tyr^_GUA_-regulated genes (Fig 5A-B). Despite these reductions at the protein level, the transcript levels of two of these three downstream genes were not significantly altered upon H_2_O_2_ treatment (Fig S5A). The finding that one downstream gene (EPCAM) is altered both at the transcript and protein levels may signify lack of translational regulation for this gene or that translational repression occurs and is accompanied by mRNA decay. Our findings are consistent with translational repression of two specific growth-regulatory genes upon tRNA^Tyr^_GUA_ depletion.

**Figure 5.**
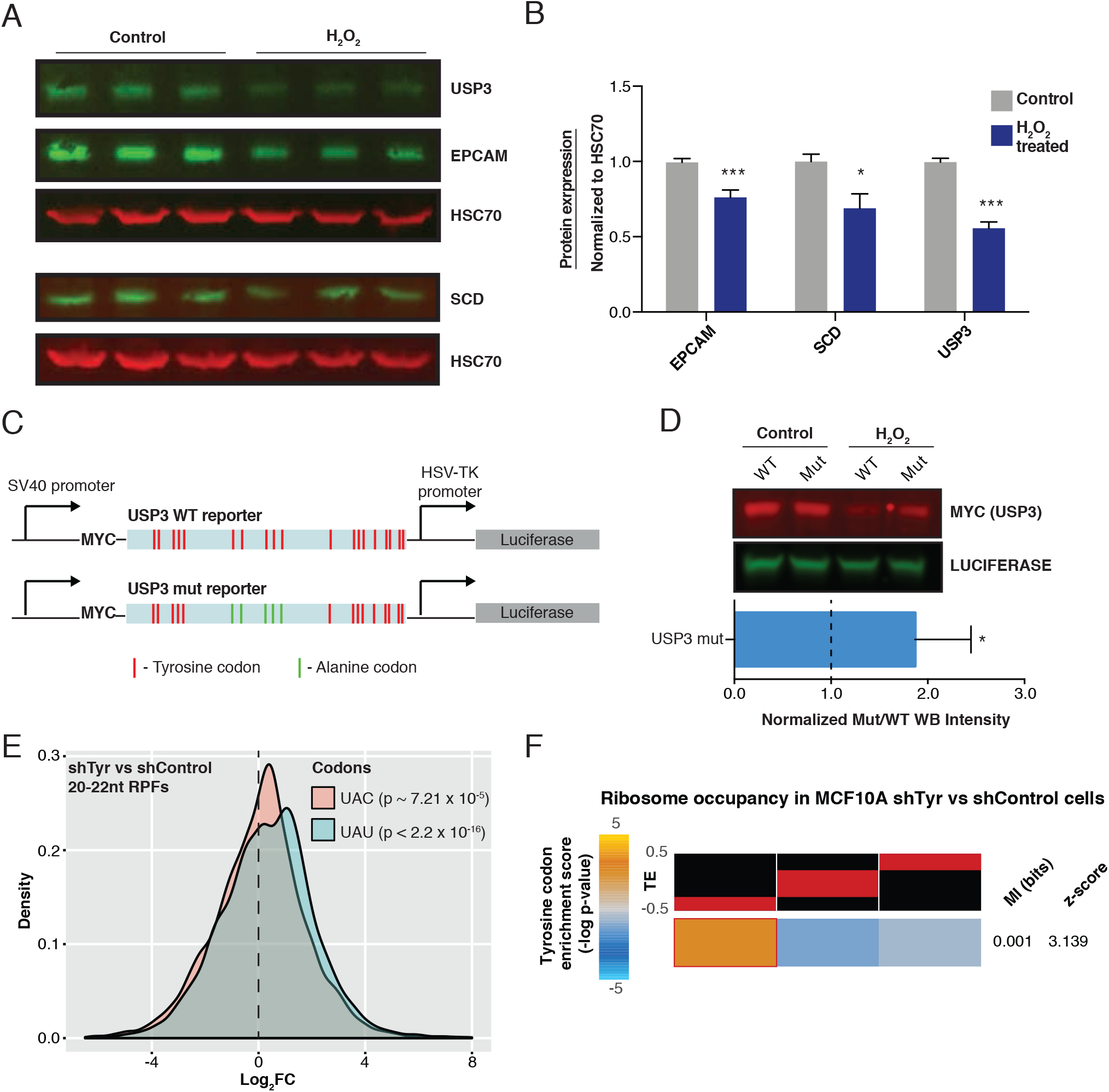
Global ribosome occupancy analysis from tRNA^Tyr^_GUA_-depleted cells reveals reduced translation efficiency for Tyr-enriched genes. (A) Quantitative western blot EPCAM, SCD, and USP3 in MCF10A cells 24 hours after treatment with H_2_O_2_ (200μM). HSC70 was used as a loading control. (B) Quantification of western results in (A) (n=9). A one-tailed Mann-Whitney test was used to establish statistical significance between treated and control conditions. (C) A schematic of the codon-based USP3 reporter. A Myc-tagged WT or mutant reporter with 5 Tyr codons mutated to Ala codons were cloned upstream of a Luciferase used for transfection normalization. (D) Quantitative western blot for the Myc-tag and Luciferase (top) with normalized fluorescent intensities (bottom) are shown (n=3). (E) Ribosome occupancy of 20-22nt RPFs in tRNA^Tyr^_GUA_-depleted cells compared to control cells reveal greater occupancy at both tyrosine codons in tRNA^Tyr^_GUA_-depleted cells. (F) Genes were sorted based on their changes in GC-corrected translation efficiency (TE) values, with reduced TE in tRNA^Tyr^_GUA_-depleted cells shown in left and enhanced TE shown on right. The red bars over each column depict the range of values in that bin. We then assessed the distribution of genes with high tyrosine codon content across these three bins using mutual information calculation and testing (see methods for details). For visualization, we used the hypergeometric distribution to assign p-values to the overlap between tyrosine-rich genes and each of the three bins. We then defined an enrichment score as –log of p-value, if there was a significant enrichment. If the overlap is significantly fewer than expected by chance, log of p-value is used instead (depletion). The resulting enrichment score is then shown as a heatmap with gold depicting positive enrichment. Data represent mean ± s.e.m. *p < 0.05, **p < 0.01, and ***p < 0.001

We took two independent approaches to test whether tRNA^Tyr^_GUA_ modulation directly impacts gene regulation at the translational level. In the first approach, we employed a codon-dependent reporter of an endogenous target of tRNA^Tyr^_GUA_ depletion. A Myc-tagged USP3 coding sequence was cloned upstream of Luciferase, which acted as a transfection control. The cloned USP3 sequence was either wild-type (WT) or a mutant variant, which contained 5 tyrosine codons mutated to alanine codons (Fig 5C). We hypothesized that if USP3 was sensitive to, and directly regulated by tRNA^Tyr^_GUA_ modulation, then reducing its tyrosine codons would reduce its susceptibility to repression at the protein level upon tRNA^Tyr^_GUA_ depletion. These reporters were transfected into MCF10A cells and 24 hours after H_2_O_2_ treatment quantitative western blotting was performed. Consistent with our model of a codon-dependent regulation at the level of translation, we noted a significant increase in the abundance of the mutant USP3 relative to the WT version of the protein (Fig 5D).

These proteins’ expression levels are sensitive to both the abundance of tRNA^Tyr^_GUA_ as well as its charging enzyme—implicating modulation of ribosomal translation of this gene set in a tRNA^Tyr^_GUA_ and tyrosine codon-dependent manner. To directly test if ribosomal engagement of tyrosine-codon enriched transcripts is impaired upon tRNA^Tyr^_GUA_ depletion, our second experimental approach was to perform ribosomal profiling in control and tRNA^Tyr^_GUA_–depleted cells (Ingolia et al., 2009). We compared the ribosome protected fragments (RPFs) detected in cells with and without tRNA^Tyr^_GUA_ depletion, in order to examine the global translational effects due to modulating this single tRNA. We observed similar length distribution and nucleotide periodicity for ribosome-protected fragments as those of previous studies (Fig S5B-C) (Ingolia et al., 2009; Lareau et al., 2014; McGlincy and Ingolia, 2017). Analysis of the 20-22nt RPFs, which represent when the A site of the ribosome is empty (Wu et al., 2019), showed an increase of reads containing either UAC or UAU codons that encode for tyrosine in tRNA^Tyr^_GUA_–depleted cells compared to control cells (Fig 5E). Consistent with previous reports for non-optimal or rare tRNAs, we observe ribosomal slowing or stalling upon decoding of tyrosine codons at the ribosomal A-site when the cognate tRNA becomes reduced (Fig S5D). Furthermore, a corrected ribosome-occupancy score was calculated for each gene as a metric for active translation in control and tRNA^Tyr^_GUA_–depleted cells. Genes with distinct translation efficiencies, defined as the ratio between RPFs and mRNA fragments, were separated into three equally populated gene sets. Genes with higher tyrosine codons were significantly enriched in the set of genes with the lowest translational efficiency (denoted by the lowest red bar) upon tRNA^Tyr^_GUA_ depletion (Fig 5F). These findings reveal that ribosomal translation of a set of tyrosine codon-enriched genes in these mammalian cells is sensitive to depletion of tRNA^Tyr^_GUA_ to physiological levels.

### Tyrosine tRF interacts with the RNA-binding proteins hnRNPA1 and SSB

In addition to causing tRNA^Tyr^_GUA_ depletion, oxidative stress also induced generation of tRF^Tyr^_GUA_ (Fig 1B, 2A, 2D). Significant tRF^Tyr^_GUA_ induction was observed at 5 minutes, remained elevated for up to 8 hours and declined to near baseline levels at 24 hours (Fig 6A-B) post H_2_O_2_ exposure. Previous studies have implicated tRFs in multiple biological processes including proliferation, cell invasion, translation, trans-generational inheritance, and cancer metastasis (Chen et al., 2016; Goodarzi et al., 2015; Honda et al., 2015; Keam et al., 2017; Kim et al., 2017; Sharma et al., 2016). Sequencing of the tRF^Tyr^_GUA_ after gel extraction identified tRFs with a 5’ leader sequence from nearly every tRNA^Tyr^_GUA_ genomic locus (Fig S6A-B). As the tRF originates from the pre-tRNA, we speculated that the tRNA splicing machinery might be involved in the induction of this fragment. Pre-tRNA processing and maturation have been well characterized (Hopper and Nostramo, 2019). In order to test this, we used RNAi-mediated knockdown for TSEN2, the catalytic subunit of the tRNA splicing complex (Paushkin et al., 2004), as well as for ANG, a RNase previously described to cleave certain tRNAs at the anticodon loop (Fu et al., 2009). Depletion of neither ANG nor TSEN2 impaired tRF^Tyr^_GUA_ formation, suggesting that these ribonucleases are not mediating this oxidative-stress induced response (Fig S6C-E). Further evidence suggesting that a defect in tRNA splicing was not the source of the tRF induction was that equal levels of tRFs were observed following RNAi-mediated depletion of the RNA kinase CLP1 (Fig S6F-G), a component of the tRNA splicing complex (Weitzer and Martinez, 2007).

**Figure 6.**
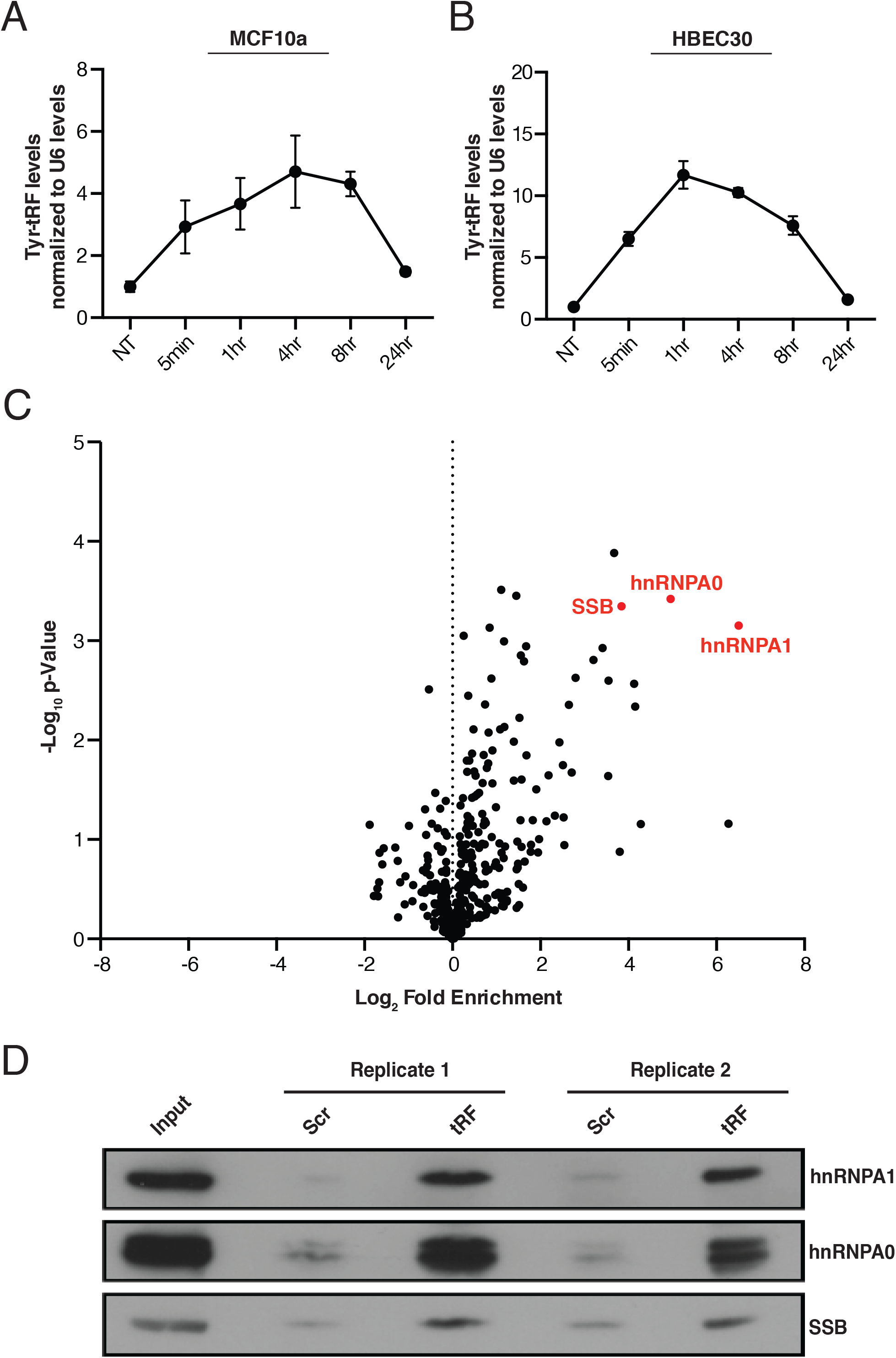
Identification of proteins that interact with tRF^Tyr^_GUA_. (A-B) Quantification of tRF^Tyr^_GUA_ induction in response to oxidative stress as a function of time in MCF10A (A) (n=4) and in HBEC30 (B) (n=6). (C) Volcano plot of mass spectrometry results from a synthetic 5’-biotinylated tRF^Tyr^_GUA_ co-precipitation experiment with cell lysate. Log2 fold enrichment values of proteins identified from tRF^Tyr^_GUA_ relative to scrambled tRF control samples. (D) Western blot validation of mass spectrometry results for three of the top hits in (A), showing co-precipitation of endogenous proteins with transfected tRF^Tyr^_GUA_ relative to a scrambled control sequence (Scr).

We next assessed whether tRF^Tyr^_GUA_ might impact the same molecular pathway as the mature tRNA. Using the conserved region from the different tRF^Tyr^_GUA_ sequences (Fig S6B), we transfected a 37nt synthetic mimetic as a means of eliciting gain-of-function. TRF^Tyr^_GUA_ transfection did not significantly alter protein levels of USP3, SCD, or EPCAM and did not impact growth (Fig S6F-G). These findings suggested that tRF^Tyr^_GUA_ might play a regulatory role independent of the tRNA^Tyr^_GUA_-mediated response. One mechanism by which tRFs have been proposed to function is through their interaction with various RNA binding proteins (RBPs) (Couvillion et al., 2010; Goodarzi et al., 2015; Haussecker et al., 2010). We hypothesized that tRF^Tyr^_GUA_ may not only be a degradation product of tRNA fragmentation, but may also interact in trans with an RBP. To test this, we used a synthetic tRF^Tyr^_GUA_ as bait in an in vitro co-precipitation experiment where the 5’-biotinylated tRF mimetic was captured on streptavidin beads and then incubated with cellular lysate. Proteins interacting with the mimetic were identified by in-solution digestion and mass-spectrometry, and compared to proteins interacting with a scrambled mimetic. Mass spectrometry identified numerous proteins that were enriched in the tRF^Tyr^_GUA_ co-precipitation relative to scrambled control oligonucleotide (Fig 6C). We selected the most significantly enriched proteins—hnRNPA0, hnRNPA1, and SSB—and validated their interaction with synthetic tRF^Tyr^_GUA_ by western blot. Western blot analyses confirmed the mass spectrometry results, revealing increased interactions between these proteins and tRF^Tyr^_GUA_ mimetic relative to scrambled control (Fig 6D). These results suggest that tRF^Tyr^_GUA_ may interact with one or more RBPs.

### Endogenous tRF^Tyr^_GUA_ interacts with endogenous hnRNPA1 and SSB

We next sought to determine if there exists an endogenous interaction between tRF^Tyr^_GUA_ and an RBP. UV-crosslinking enables assessment of direct endogenous interactions between RBPs and their cellular RNA substrates (Mili and Steitz, 2004; Ule et al., 2003) and has been coupled with deep-sequencing methods such as HITS-CLIP or PAR-CLIP to identify the landscape of RNAs that interact with a given RBP (Hafner et al., 2010; Licatalosi et al., 2008). The initial steps of such CLIP-sequencing approaches require limited RNase digestion of immunoprecipitated ribonucleoprotein complexes prior to SDS-PAGE visualization. Such experiments have previously been done with Argonaute-2, which binds microRNAs (Chi et al., 2009), and YBX1, which binds tRFs (Goodarzi et al., 2015). These experiments have revealed well-defined bands roughly the size of the RBP, which represent the RBP bound to a population of small RNAs. In contrast, for RBPs that bind mRNAs, CLIP-seq methods reveal a smear representing the RBP bound to a population of such longer RNAs (Chi et al., 2009; Goodarzi et al., 2015) (Fig 7A). To determine if any of the candidate RBPs identified by mass spectrometry interact with endogenous small RNA populations, we included an experimental condition where the HITS-CLIP protocol was conducted in the absence of RNase digestion to ensure that any potential small RNA-RBP bands visualized were not a consequence of, or confounded by, RNase digestion. UV-crosslinked immunoprecipitation followed by SDS-PAGE in the presence or absence of RNase digestion revealed that endogenous hnRNPA1 and SSB interacted with an endogenous small RNA population (Fig S7A-B). We did not observe a small RNA-ribonucleoprotein band for hnRNPA0 in the absence of RNase (Fig S7C), even upon prolonged autoradiographic exposure, suggesting that this RBP either does not significantly interact with a small RNA population *in vivo* or this method is not conducive to identifying this interaction. These findings suggest that hnRNPA1 and SSB directly interact with small RNA populations *in vivo*.

**Figure 7.**
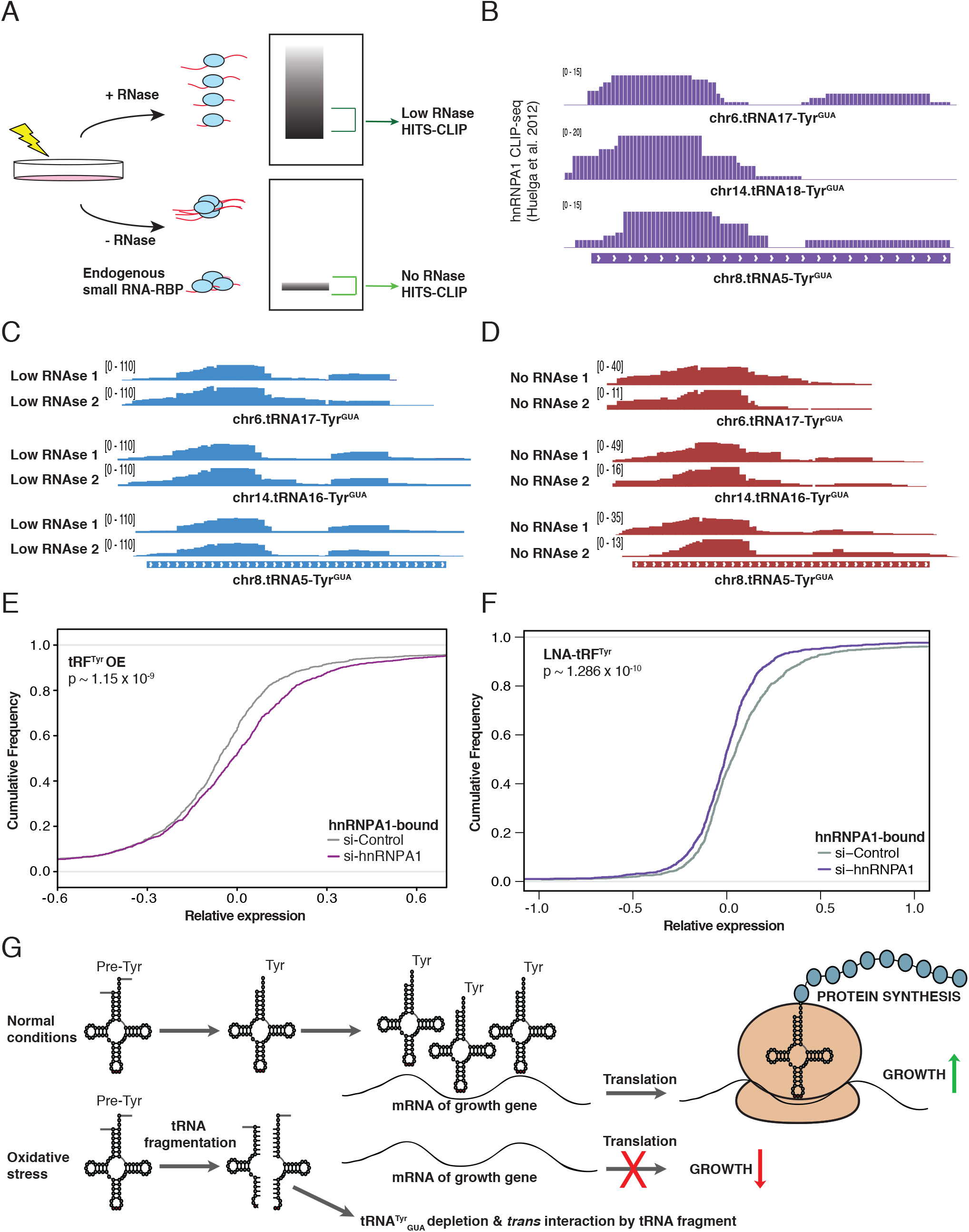
HITS-CLIP reveals endogenous SSB binding to endogenous tRF^Tyr^_GUA_. (A) Schematic depicting the expected visualization of a cross-linked immunoprecipitation by autoradiogram. Samples processed in the absence of RNase digestion that revealed a band corresponding to smRNA-RBP interactions were processed for HITS-CLIP. Samples processed using low RNase digestion showing a smear representing mRNA-RBP interactions by autoradiogram were also processed for HITS-CLIP. (B) IGV plots from the hnRNPA1 HITS-CLIP (Huelga et al., 2012) reveals interactions with the tRF^Tyr^_GUA_. (C) IGV plots representing SSB interacting with the tRF^Tyr^_GUA_ in samples that were treated with low levels of RNase digestion. SSB bound tRF^Tyr^_GUA_ reads mapped to multiple loci encoding tRNA^Tyr^_GUA_. (D) Similar to the IGV plots shown in (C), but depicting SSB interactions with tRF^Tyr^_GUA_ loci in samples without RNase digestion. (E) A cumulative distribution in control and hnRNPA1 depleted cells of the expression levels for mRNA transcripts with 3’ UTR hnRNPA1 CLIP binding (Huelga et al., 2012). Transfection of tRF^Tyr^_GUA_ led to a significant right-shift in the expression levels of 3’ UTR bound hnRNPA1 transcripts. Statistical significance was measured using the Kolmogorov-Smirnov test. (F) Cumulative distribution as in (E). Transfection of locked nucleic acid against tRF^Tyr^_GUA_ and treatment with 200µM H_2_O_2_ led to a significant left-shift in mRNA stability of 3’UTR bound hnRNPA1 transcripts. Statistical significance was assessed using the Kolmogorov-Smirnov test. (G) Model of tRNA^Tyr^_GUA_-dependent gene regulatory response to oxidative stress.

We next sought to define the identities of the small RNAs bound by hnRNPA1 and SSB. An analysis of tRFs bound in a previously published CLIP-seq study for hnRNPA1 (Huelga et al., 2012) validated our observations by revealing a reciprocal interaction between hnRNPA1 and tRF^Tyr^_GUA_ (Fig 7B). As an abundant RBP, hnRNPA1 has many previously described roles in gene expression regulation, including as a regulator of splicing and mRNA stability via binding to 3’ UTRs (Glisovic et al., 2003; Wang et al., 2016). HnRNPA1 has also been implicated in promoting growth and multiple cancer progression phenotypes (Biamonti et al., 1993; Roy et al., 2017). We next conducted HITS-CLIP for endogenous SSB with and without RNase digestion. SSB, also known as La, is a well characterized RNA-binding protein known to bind the nascent 3’ ends of Pol III transcripts, including those of full-length pre-tRNAs (Gottlieb and Steitz, 1989; Maraia et al., 1994; Yoo and Wolin, 1997). Beyond its described nuclear roles in Pol III transcript binding, key cytoplasmic functions for SSB have also been reported including regulating translation of certain cellular and viral mRNAs (Costa-Mattioli et al., 2004; Maraia et al., 2017). Although our HITS-CLIP experiment was designed to enrich for signal from tRFs, we found that consistent with its previously described canonical role in binding Pol III transcripts, alignment and analysis of sequencing reads revealed binding of SSB to the 3’-trailers of pre-tRNAs (Fig S7D). Importantly, in addition to this previously described binding, we observed previously unreported interactions of SSB with the 5’ half of tRNA^Tyr^_GUA_ arising from multiple distinct loci (Fig 7C). In experiments with and without RNase, sequencing reads containing and not containing the 5’ leader region of pre-tRNA^Tyr^_GUA_ were detected. The 5’ leader containing reads likely represent intermediates in the pre-tRNA^Tyr^_GUA_ processing reaction (Fig 7C-D) (Hanada et al., 2013). The abundant number of reads mapping to the 5’ regions of tRNA^Tyr^_GUA_ distinguish these SSB-tRF interactions from the previously described canonical SSB interactions with 3’-trailers of full-length pre-tRNAs. This notion is further supported by our observations of SSB binding to the 5’ tRF without 3’-trailer binding even in the absence of RNase digestion. These observations reveal that tRF^Tyr^_GUA_ interacts in trans with RBPs in human cells.

We next investigated if the stress-induced tRF^Tyr^_GUA_ might regulate the activity of hnRNPA1 or SSB as a trans factor. Using the 3’ UTR targets of hnRNPA1 previously identified by CLIP-seq (Huelga et al., 2012), we assessed whether tRF^Tyr^_GUA_ impacts hnRNPA1-mediated mRNA stabilization. Following transfection of the tRF^Tyr^_GUA_ mimetic or a scrambled control, we observed impaired stabilization of hnRNPA1 targets in an RBP-specific manner (Fig 7E). Conversely, locked nucleic acid mediated inhibition of tRF^Tyr^_GUA_ or a control had the opposite effect in increasing stability of hnRNPA1 mRNA targets in an hnRNPA1 specific manner (Fig 7F). These results are consistent with known reports of hnRNPA1 binding to 3’ UTRs to increase mRNA stability (Glisovic et al., 2003; Wang et al., 2016). Taken together, our results are consistent with tRF^Tyr^_GUA_ competing with hnRNPA1 target transcripts for binding to endogenous hnRNPA1. TRF^Tyr^_GUA_ -dependent reduction of hnRNPA1 binding to it’s 3’ UTR targets decreased stability of hnRNPA1 transcripts during α-amanitin-stability analyses. We next sought to identify potential effects of tRF^Tyr^_GUA_ on SSB function. Transfection of the tRF^Tyr^_GUA_ mimetic did not impact expression of Pol III transcriptional targets, suggesting that the canonical nuclear role for SSB was unaffected by increased levels of the tRF^Tyr^_GUA_ mimetic in this context (Fig S7E). Our findings describe that stress-induced fragmentation can cause a specific transfer RNA to become depleted, resulting in translational consequences, and also give rise to a tRNA fragment that can interact in trans and functionally impact the regulon of an RNA binding protein known to promote growth and cancer progression phenotypes.

### TRNA^Tyr^_GUA_ fragment generation is DIS3L2 exoribonuclease dependent

We next sought to identify a ribonuclease involved in oxidative-stress induced tRF^Tyr^_GUA_ generation. In an ongoing RNAi screen focused on identifying putative regulators of tRNA fragments in *C*.*elegans*, we observed that depletion of the exoribonuclease *disl2*—the ortholog of *DIS3L2*—a tumor suppressor in Wilms tumor, impaired stress-induced tRNA fragment levels (unpublished results). Exposure of worms to H_2_O_2_ for a brief period (15 minutes) so as to avoid starvation effects revealed *disl2*-depleted animals to produce less stress-induced tRNA^Tyr^_GUA_ fragments relative to control animals (Fig S7F-H). A similar effect was observed in *disl2* mutant (*syb1033*) worms upon oxidative stress exposure. Depletion or genetic deletion of DIS3L2 also impaired, but did not completely eliminate, oxidative stress-induced tRNA^Tyr^_GUA_ fragment levels in human cells (Fig S7I-J). DIS3L2 depletion selectively impaired tRF^Tyr^_GUA_ formation upon oxidative stress, and did not cause pre-tRNA^Tyr^_GUA_ accumulation— consistent with DIS3L2 mediating processing/maturation of a tRF^Tyr^_GUA_ precursor rather than the pre-tRNA^Tyr^_GUA_ cleavage. These findings implicate the DIS3L2 tumor suppressor in tRF^Tyr^_GUA_ generation.

## DISCUSSION

Aberrant tRNA breakdown products were first detected nearly forty years ago upon analyses of the urine of cancer patients (Gehrke et al., 1979). The biological and molecular roles of such small RNAs, commonly termed tRFs, have recently received considerable attention. TRFs have been implicated in processes such as hematopoiesis (Guzzi et al., 2018), cancer pathogenesis (Goodarzi et al., 2015; Lee et al., 2009), and shown to mediate effects via interactions with RNA-binding proteins (Couvillion et al., 2010; Goodarzi et al., 2015) or messenger RNAs (Kim et al., 2017). Early studies of specific tRFs in *Saccharomyces cerevisiae* revealed that the specific tRNA pools studied were not impacted upon tRNA fragmentation (Saikia et al., 2012; Thompson and Parker, 2009b). Similar observations were made in mammalian systems for specific tRFs (Yamasaki et al., 2009). By comprehensively surveying global tRF generation and tRNA abundances, we have identified a specific tRNA, tRNA^Tyr^_GUA_, for which significant depletion occurs during stress-induced fragmentation. Such stress-induced depletion resulted in functional consequences of tyrosine codon-dependent translational repression and growth suppression.

The advent of numerous methods that enabled profiling of tRNAs revealed widespread alterations in tRNA expression levels in cancer (Gingold et al., 2014; Goodarzi et al., 2016; Pavon-Eternod et al., 2009). Genomic copy number gains at tRNA loci have been shown to account for enhanced expression of specific tRNAs that functionally drive cancer progression (Goodarzi et al., 2016; Truitt and Ruggero, 2016). This has raised the question of whether in the absence of genomic instability, endogenous pathways exist for physiological modulation of tRNA levels with functional consequences. Our observations reveal that in non-malignant epithelial cells, there exists an oxidative stress response pathway that represses growth upon selective tRNA depletion. The consequence of this response is repressed translation of a set of growth promoting genes. This tRNA depletion response likely cooperates with other cellular oxidative response pathways (Martindale and Holbrook, 2002). While our work has interrogated the consequences of tRNA^Tyr^_GUA_ depletion, future work is needed to determine the impact, if any, of depletions of other tRNAs that were observed to generate fragments and also become depleted. Two such examples are tRNA^Leu^_UAA_ and tRNA^Leu^_CAG_. Interestingly, although there exist six isoacceptor tRNAs for leucine, we only observed significant concomitant tRF induction and tRNA depletion for these two isoacceptors, suggesting potential tRNA fragmentation selectivity amongst the isoacceptors for a given amino acid, with potential for codon-biased translational consequences.

Previous studies have assessed the relationship between global tRNA modulations and effects on translation in *E. coli* (Wohlgemuth et al., 2013; Zhong et al., 2015). Consistent with our findings, the studies in *E. coli* have revealed that protein translation can be impacted by cellular tRNA availability. While oxidative stress has been observed to repress global tRNA levels in *E. coli* (Zhong et al., 2015) and *S. cerevisiae* (Torrent et al., 2018), our results reveal that in mammalian cells, there is a selective tRNA and codon-dependent response to oxidative stress. Differences in stress response between mammalian cells and unicellular organisms suggests that higher organisms may have evolved mechanisms for selective tRNA modulation in response to a key cellular stress. Future studies are warranted to investigate these possibilities. While we do not yet know the mechanistic basis for this selectivity, we speculate that the tRNAs that become modulated upon oxidative stress may be of lower relative abundance compared to other unaffected tRNAs or that various steps in tRNA processing might be targets of regulation. In humans, all tRNA^Tyr^_GUA_ isodecoder genes have introns that require splicing to generate mature tRNAs. Our observed tRNA^Tyr^_GUA_-dependent response initiates at the pre-tRNA^Tyr^_GUA_ level, giving rise to tRFs, and consequently reduces the abundance of mature tRNA^Tyr^_GUA_.

Further investigation into tRNA processing may elucidate additional mechanisms by which cells post-transcriptionally regulate protein expression. Though our data indicates that tRNA splicing is not the main source of tRF induction, we cannot rule out the possibility that additional currently unknown RNases might also have a role in the tRNA splicing complex. If the known tRNA splicing complex is involved, conditional genetic inactivation of components of the tRNA splicing complex or TSEN2 might be necessary to completely blunt tRF induction upon oxidative stress. Dysregulated tRNA splicing has been previously implicated in human disease. Aberrantly generated tRFs derived from pre-tRNA introns have been linked with human neuronal degeneration (Karaca et al., 2014; Schaffer et al., 2014). Furthermore, by reacting with nucleic acids, proteins, and lipids, reactive oxygen species can cause global cellular dysfunction (Cross et al., 1987; Radi, 2018). Consistent with this, oxidative stress has been implicated or associated with a myriad of human diseases including Parkinson’s, ALS, Alzheimer’s, cardiovascular disease, and cancer pathogenesis (Cervantes Gracia et al., 2017; Lin and Beal, 2006; Piskounova et al., 2015). Exposure of mouse neurons and MEFs to oxidative stress was previously shown to elicit fragmentation of tRNAs, including tyrosine tRNA (Hanada et al., 2013). Sequencing analysis revealed generation of fragments from tyrosine pre-tRNA—similar to our observations (Fig 1a). This past work, however, did not assess the subsequent impact of tRF formation on pre-tRNA or tRNA abundances in the neural cells under study. It would be of interest to determine if oxidative stress-induced fragmentation could also functionally deplete the corresponding tRNA pool in non-mitotic neurons in a manner similar to that we observed in mitotic epithelial cells in this study.

In addition to the depletion of tRNA^Tyr^_GUA_ in response to oxidative stress, we have implicated hnRNPA1 and SSB/La as RNA binding proteins with direct interactions with tRF^Tyr^_GUA_. SSB/La has numerous cellular functions, perhaps chief amongst them being pre-tRNA processing (Gottlieb and Steitz, 1989; Wolin and Cedervall, 2002). While future work is necessary to define the potential regulatory role of this interaction, the interaction of a stress-induced tRNA fragment with a central regulator of physiological tRNA processing suggests a potential regulatory feedback loop that may impact tRNA maturation upon stress induction. Similar to SSB, hnRNPA1 has multiple known roles in regulating RNAs (Jean-Philippe et al., 2013), such as binding to 3’ UTRS to increase mRNA stability (Glisovic et al., 2003; Wang et al., 2016). Our data suggests that the tRF^Tyr^_GUA_ competes with these 3’ UTRs for binding to hnRNPA1 and as a consequence, decreases the stability of these hnRNPA1 targets. Interestingly, past studies have also implicated tRFs in impairing stability and expression of translation genes (Goodarzi et al., 2015) and altering the expression of a ribosomal transcript (Kim et al., 2017). Our study explores a divergent response to oxidative stress that surprisingly originates from a tRNA precursor. The rapidly induced tRF^Tyr^_GUA_ acts to destabilize mRNAs in an hnRNPA1-dependent manner while the delayed tRNA^Tyr^_GUA_ reduction mediates translational repression and growth suppression. These studies in total suggest a natural link between tRFs and the central cellular process in which their precursor molecules (tRNAs) participate.

Our findings raise a number of questions for future study. Firstly, could tRNA^Tyr^_GUA_ fragmentation and depletion occur during normal development and physiology to mediate growth repression? Oxidative species are generated during cellular respiration and may endogenously activate this response. Moreover, immune cells generate free radicals as a means of combatting pathogens (Dahlgren and Karlsson, 1999; Radi, 2018). Such processes may modulate tRNA^Tyr^_GUA_ levels and elicit growth phenotypes. Independent of oxidative stress, tRNA^Tyr^_GUA_ fragmentation and depletion may be regulated via other stresses, signaling pathways, developmental events or in the context of normal physiology. Developmental surveys of tRF^Tyr^_GUA_ or tRNA^Tyr^_GUA_ levels in diverse tissues may shed light on these possibilities. Moreover, our identification of the growth-associated gene *DIS3L2* as an exoribonuclease involved in tRNA^Tyr^_GUA_ fragmentation across multiple species suggests that this tRNA depletion response may perhaps also play a role in these other contexts. Recent elegant genetic studies revealed that the previously assumed direct substrate of DIS3L2 is not let-7, suggesting that another RNA substrate may mediate the effects of DIS3L2 on tumorigenesis and growth (Hunter et al., 2018). Future work is needed to interrogate the role of this tRNA fragmentation response in the pathogenesis of this disorder. More broadly, our studies of tRNA modulation by oxidative stress raise the possibility that additional stresses may generate similar or distinct tRNA fragmentation and depletion effects, which may mediate coherent gene-regulatory responses. The degeneracy of the genetic code coupled with the wide spectrum of metabolic modifications on tRNAs positions tRNAs as ideal molecules for ‘sensing’ metabolic derangements and mediating rapid transcription-independent gene expression responses.

## EXPERIMENTAL PROCEDURES

### Cell culture

MCF10A cells were cultured in DMEM/F12 media supplemented with 5% horse serum and final concentrations of 20 ng/ml of EGF, 0.5 mg/ml of hydrocortisone, 100 ng/ml of cholera toxin, and 10 μg/ml of insulin. HBEC30 cells were cultured in keratinocyte-SFM media with the included supplements of BPE and EGF. All cell lines were STR tested and routinely tested for mycoplasma contamination. To induce oxidative stress, cells were incubated with 200μM H_2_O_2_ or 50μM of menadione (Sigma) dissolved in DMSO in their respective medias for the time specified in each experiment. RNA was then isolated by TRIzol and isopropanol precipitation as described below.

### Stable cell line generation

Lentivirus was produced in 293T cells grown in 10 cm plates. 3 μg of each packaging vector (pRSV-Rev, pCMV-VSVG-G, and pCgpV) were transfected with 9 μg of the appropriate hairpin in a pLKO-backbone vector using 30 μl of Lipofectamine 2000 (Invitrogen). After 24 hours, the media was replaced with fresh media and virus-containing supernatant was harvested 48 hours after transfection. The supernatant was filtered through a 0.45 μm filter before 2 ml of virus was used to transduce MCF10A cells plated in 6-well plates with 8 μg/ml of polybrene. Media was changed after 24 hours and antibiotic selection with 1 μg/ml of puromycin was started 48 hours after virus transduction. Targeted sequences used for stable RNAi cell lines were: TTATGCCACAGTCCTTATAAT (YARS sh2), GTTCATCAAAGGCACTGATTA (YARS sh3), and TGGTAGAGCGGAGGACTGTAGA (tRNA^Tyr^_GUA_). Cells were selected for 3-5 days with a population of non-transduced cells as a control. Knockdown of mRNA and protein were validated by qRT-PCR and western blot, respectively, while knockdown of tRNA was validated by northern blot.

### RNA isolation and purification

RNA was extracted from cells using TRIzol and isopropanol precipitation at -20°C overnight. After centrifugation at max speed (∼21,000 x g) in a refrigerated tabletop centrifuge, the RNA pellet was washed twice with ice-cold 75% EtOH before being resuspended in RNase free water or TE-buffer.

### Northern blotting

Purified RNA was run on a 10% Urea-PAGE gel before being transferred onto a nylon membrane and UV crosslinked (240 mJ/cm^2^). The membrane was pre-hybridized in UltraHyb-Oligo buffer (Ambion) at 42°C. DNA oligos were radiolabeled with [γ-^32^P]ATP using T4 PNK (NEB) and further purified by G-25/G-50 columns before incubating with the blot overnight. After hybridization, the blot was washed twice with SSC and SDS buffers before being developed. Probes that were ^32^P labeled and used for detection were: ACAGTCCTCCGCTCTACCAGCTGA (tRNA^Tyr^_GUA_), ACAGTCCTCCGCTCTACCAACTGA (*C. elegans* tRNA^Tyr^_GUA_), CAGCGCCTTAGACCGCTCGGCCA (tRNA^Leu^_HAG_), AACGCAGAGTACTAACCACTATACG (tRNA^His^_GUG),_ GCGCCGAATCCTAACCACTAGACCA (tRNA^Glu^_YUC_), CACGAATTTGCGTGTCATCCTT (U6), and CAAATTATGCAGTCGAGTTTCCCACATTTG (U1). Quantification was done using FIJI (ImageJ) where the intensity of each band over background was measured and normalized to U6 levels.

### Quantitative LC-MS/MS proteomic profiling

For label-free quantitation analysis of protein levels by mass spectrometry, tRNA^Tyr^_GUA_-depleted and YARS-depleted cells were compared control hairpin expressing cells. For each set of samples, 50ug of lysate (n=3 per condition) was ice-cold acetone precipitated. Precipitates were dissolved in 50uL 8M Urea/0.1M ammonium bicarbonate/10mM DTT. After 45 minutes of incubation at room temperature, reduced cysteines were alkylated with iodoacetamide (Sigma) in the dark for 45 minutes. Volumes were diluted 2-fold and proteins were digested overnight with 1 µg Lysyl Endopeptidase (Wako). Prior to trypsinization (1ug) (Promega), the samples were diluted an additional 2-fold (0.1 M ammonium bicarbonate). After 8h, the digestions were halted by the addition of neat TFA (Sigma) and then peptides were desalted and concentrated using stage tips.

Peptides were separated using a direct-loading setup with a 50cm EasySprayer C_18_ column (ES80, Thermo). Peptides were eluted using a gradient increasing from 2% B/98% A to 45% B/55% A (A: 0.1% formic acid, B: 80% acetonitrile/0.1% formic acid) in 220 minutes. The gradient was delivered at 300nL/min (Easy 1200, Thermofisher). The mass spectrometer (Fusion Lumos, Thermo Fisher) was operated in High/High mode with MS and MS/MS mass resolution being 120,000 and 30,000, respectively. For MS/MS, Automatic Gain Control was set to 1e5 with a maximum injection time of 54 ms. MS-data were queried against Uniprot’s Human Complete Proteome (70,246 sequences, March 2016) using MaxQuant v1.6.0.13. Oxidized methionine and protein N-terminal acetylation were allowed as variable modifications and up to 2 missed cleavages were allowed. Proteins were quantitated using MaxQuant’s ‘Label Free Quantitation’ (LFQ) feature and signals were required in minimum 2-of-3 replicates for at least one condition (4,753 proteins quantitated). Missing LFQ values where imputed. All data analysis was carried out using Perseus v1.6.0.7.

### Western blotting

Cells were lysed in ice-cold RIPA buffer containing a protease inhibitor cocktail (Roche) before cellular debris was cleared by centrifugation at max speed in a refrigerated tabletop centrifuge. Samples were heated with LDS buffer and reducing agent before running on an SDS-PAGE gel and transferred onto a PVDF membrane (Bio-Rad). The membranes were blocked and then probed using target-specific antibodies. Antibodies used were YARS (Abcam, ab150429, RRID: AB_2744675, 1:1000), EPCAM (Proteintech, 21050-1-AP, RRID: AB_10693684, 1:1000), SCD (Proteintech, 23393-1-AP, RRID: AB_2744674, 1:1000) USP3 (Proteintech, 12490-1-AP, RRID: AB_10639042, 1:1000), SSB (MBL, RN074PW, RRID: AB_11124309, 1:1000), hnRNPA1 (Santa Cruz, sc-32301, RRID: AB_627729, 1:1000), hnRNPA0 (Bethyl Laboratories, A303-941A, RRID: AB_2620290, 1:1000), DIS3L2 (Novus biologicals, NBP2-38264), Myc (Cell Signaling, 2276S, RRID: AB_331783), and Luciferase (Proteintech, 27986-1-AP, RRID:AB_2750646). Loading controls used were HSC70 (Santa Cruz, sc-7298, RRID: AB_627761, 1:2000). Chemiluminescent signal was detected by HRP-conjugated secondary antibodies, ECL western blotting substrate (Pierce), and the SRX-101A (Konica Minolta) developer according to manufacturer’s instructions. Membranes were stripped (Restore western blot stripping buffer, Pierce), blocked, re-probed, and re-developed if necessary.

### Quantitative western blotting

To compare protein expression levels by western blot, the Odyssey® quantitative western blotting system (LI-COR) was used. This method is identical to western blotting except membranes were blocked in Odyssey® blocking buffer (PBS) and the secondary antibody used was a species-specific fluorescent IRDye®. Membranes were then imaged using the Odyssey® Sa Infrared Imaging System at the Rockefeller University Center for High Throughput Screening. Image quantification was done using Image Studio™ Lite and an unpaired t-test was used to determine statistical significance.

### Codon reporter assays

A Myc-tagged construct was synthesized by Genewiz and cloned into the psiCHECK2 vector (Promega), replacing the synthetic Renilla Luciferase gene. The construct contained a 6x-glycine linker was placed between the Myc-tag and the gene. The constructs included a wild-type USP3 (NM_006537) or a USP3 gene with a set of 5 tyrosines mutated to alanines. Multiple mutant constructs were cloned but only those with high levels of expression were chosen for testing. MCF10A cells were seeded to be 70% confluent in a 6 cm plate the next morning. Cells were exposed to 200μM H_2_O_2_ or a control for one hour before being transfected with the reporters using Lipofectamine 3000 (Invitrogen). Cells were lysed and compared through quantitative western blotting 24 hours post H_2_O_2_ treatment.

### Quantitative RT-PCR

To measure mRNA transcript levels, RNA was converted to cDNA (SuperScript III, Life Technologies) followed by Fast SYBR™ Green quantification (Life Technologies) according to manufacturer’s instructions. Primers used for qRT-PCR were: YARS: CTGCACCTTATCACCCGGAAC, TCCGCAAACAGAATTGTTACCT, EPCAM: AATCGTCAATGCCAGTGTACTT, TCTCATCGCAGTCAGGATCATAA, SCD: TCTAGCTCCTATACCACCACCA, TCGTCTCCAACTTATCTCCTCC, USP3: CAAGCTGGGACTGGTACAGAA, GCAGTGGTGCTTCCATTTACTT, ANG: CTGGGCGTTTTGTTGTTGGTC, GGTTTGGCATCATAGTGCTGG, TSEN2: ATGCGGAGGACATTGAGCAG, CGGTTTCGTGTAATCCTTGAGG, CLP1: GGAGTTGTTGACTGGCATGG, CCGCTCAGTTGCACAGAAC, HPRT: GACCAGTCAACAGGGGACAT, CCTGACCAAGGAAAGCAAAG, *C. elegans Y45F10D*.4: GCGAAAACACTCCTGCAC, TTTCGCGGGTTCTCGTAGTG, *C. elegans disl-2*: GTGCTGCCGCTACAGTCAA, GAACCGTCTTCGATTCCGG, *C. elegans disl-2(syb1033) ko:* GATTTTACCCGACAGCGA, TTACAGCCACCACATCTCCT. Expression levels of mRNA were performed with either an ABI Prism 7900HT Real-Time PCR System (Applied Biosystems) or a QuantStudio 5 Real-Time PCR System (Applied Biosystems).

### Small RNA sequencing

MCF10A cells were seeded at 50% confluency in duplicate into 10 cm plates. The next day, cells were exposed to 200μM H_2_O_2_ for one hour before harvesting for RNA purification. Small RNA was purified using the microRNA purification kit (Norgen Biotek Corp) according to manufacturer’s instructions. To identify the tRF^Tyr^_GUA_ from the pre-tRNA^Tyr^_GUA_, RNA was gel extracted between 40-50nt on a TBE-Urea PAGE gel. The RNA was then treated with polyphosphatase (Lucigen) before acid phenol chloroform extraction and RNA precipitation. Small RNA sequencing was performed using the NEXTflex→ Small RNA Sequencing Kit v3 (BiooScientific). Different samples were barcoded before being sequenced on the HiSeq 2000 Illumina sequencer. For computational analysis, fastq files were aligned to hg19 using bowtie2, and reads were further sorted, indexed, and counts were generated with samtools. Raw counts were imported into RStudio V1.1.383 and differential analysis was performed using DESeq2.

### Transient overexpression of tRNA^Tyr^_GUA_

Overexpression constructs for tRNA^Tyr^_GUA_ were made as previously reported by (Mattijssen et al., 2017). Briefly, 3 copies of tRNA^Tyr^_GUA_, per plasmid, were synthesized by Genewiz into the pUC57-KAN vector. Each copy had 150nt upstream and 90nt downstream genome sequences included but did not include the intron sequence. This vector or an empty pUC57-KAN vector were transiently transfected using Lipofectamine 3000 (Invitrogen).

### Cell growth assays

300,000 or 400,000 cells were seeded into two 6-well plates. Cells were trypsinized and viable cells were counted using a trypan blue and a hemocytometer on days 1 and 3 after seeding. Each experiment included three technical triplicates and was repeated three times. Two-way ANOVA was used to test for significance.

For growth assays with siRNA-mediated gene knockdown, cells were reverse transfected with siRNA (IDT) and Lipofectamine RNAimax (Invitrogen) so that they would be 70-80% confluent in a 10 cm plate. The following day, cells were seeded at a density of 200,000 cells per 6-well plate and counted on days 1 and 3 after seeding into 6-well plates. As before, two-way ANOVA was used to test for significance.

### Cell viability assays

MCF10A cells were seeded into 6-well plates and treated with 200μM H_2_O_2_ for one hour. This media was retained while cells were trypsinized and then resuspended in the previous media that would contain any cells that had died prior to trypsinization. Cell viability was measured using trypan blue exclusion and a hemocytometer. Viability was determined by (Total number of cells counted – Number of blue (nonviable) cells) / Total number of cells counted. Each experiment included three technical triplicates and was repeated twice. An unpaired t-test was used to test for significance.

### Transfer RNA profiling

MCF10A cells were cultured in duplicates to be 50% confluent in 10 cm plates. Cells were then exposed to 200μM H_2_O_2_ for 8 hours or 24 hours before RNA was isolated using the mirVana miRNA isolation kit. Transfer RNA profiling was done as described by (Goodarzi et al., 2016). Briefly, biotinylated probe-pairs against nuclear encoded tRNAs were designed to hybridize to each half of the tRNA. A nick at the anticodon end of the DNA-RNA hybrid was filled using T4 DNA ligase and SplintR ligase (NEB). MyOne-C1 Streptavidin Dynabeads (Invitrogen) were used to purify the DNA-RNA hybrids and ligated probes were eluted following RNase H and RNase A incubation. Eluted probes were PCR amplified and high-throughput sequenced.

For computational analysis, fastq files were aligned to tRNA probe sequences using bowtie2, and reads were further sorted, indexed, and counts were generated with samtools. Raw counts were imported into R V3.4.1 and normalized with EdgeR. Linear regression tests were used to assess tRNAs that were significantly depleted at 8 and 24 hour conditions relative to control conditions.

### Whole-genome ribosomal occupancy profiling

This procedure was conducted as described by (McGlincy and Ingolia, 2017). Briefly, cells were washed and flash frozen with liquid N_2_ before being lysed with lysis buffer containing cycloheximide (Alfa Aesar). Lysate was digested with RNase I (Lucigen) before ribosomes were isolated through ultracentrifugation through a sucrose cushion. The ribosome pellet was resuspended in a solubilization buffer containing 0.5% SDS and 1mM EDTA and TRIzol before RNA was extracted using the Direct-zol kit (Zymo Research). RNA was separated on a 15% TBE-Urea gel before RNA between 17nt and 34nt were gel extracted. Barcoded pre-adenylated linkers were ligated using T4 RNA ligase 2 truncated K227Q (NEB) and rRNA was depleted using the Ribo-Zero gold kit (Illumina) according to the manufacturer’s protocol. RNA was converted to cDNA using SuperScript III (Life Technologies) and the RT product was circularized by CircLigase II (Epicentre). A PCR library was amplified and sequenced using Illumina Nextseq 500 at the Rockefeller University Genomics Center.

For the ribosome footprinting data, reads were first subjected to linker removal and quality trimming (cutadapt v1.17). The reads were then distributed among the samples based on their assigned barcodes using fastx_barcode_splitter (using --eol and --mismatches 1). The reads were then collapsed and UMIs were extracted in two steps (2 at the 5’ end and 5 at the 3’ end) using UMI Tools. The reads were then aligned against a reference database of rRNAs and tRNAs as to remove contaminants (using bowtie 2.3.4.1). STAR was then used to align the remaining reads to the human transcriptome (build hg38). PCR duplicates were then removed using UMI Tools. Xtail (Xiao et al., 2016) was used to count RPFs, estimate translation efficiency, and perform statistical comparisons. For RNA-seq data analysis, reads were first subjected to quality trimming and adapter removal. STAR (v2.5.2a) was used to align the reads to the human transcriptome (hg38). The number of reads mapping to each gene was counted using htseq-count.

### Analysis of translational efficiency from ribosome profiling

The logTER (log2 of translation efficiency ratio) between shTyr and shControl samples calculated by Xtail showed a substantial and significant GC bias. As such, prior to further analysis, we first corrected the logTER values by fitting a linear model based on GC content of each genes as the co-variate. We then used the GC-independent residual values for further analysis. We asked whether stratifying genes based on their Tyr content is informative of corrected TER measurements. For this, we performed a t-test between two groups defined by the Tyr content of the 75^th^ percentile. At this threshold, we observed a significantly lower TE for in the shTyr samples (P<0.0002). To perform a confirmatory analysis, we also performed a gene-set analysis using our iPAGE platform (Goodarzi et al., 2009). This analysis also revealed a significant enrichment of genes with high Tyr content (same threshold as above) among those in the bottom tertile of the GC-corrected logTER values.

### Using the tRF^Tyr^_GUA_ mimetic to identify interacting proteins

A 5’ biotinylated 37-nucleotide tRF^Tyr^_GUA_ mimetic (CCUUCGAUAGCUCAGCTGGUAGAGCGGAGGACUGUAG) and a control oligo (GAGACCAGGGUACGCAAUCGAGUUGUUGGGCACUCUG) that was a scrambled version of the tRF^Tyr^_GUA_ were synthesized (IDT). Each mimetic was incubated at 4°C with equal amounts of MCF10A cell lysate containing protease inhibitor that had been pre-cleared for debris and by the control oligo. Proteins that bound to our mimetic were co-precipitated by MyOne-C1 Streptavidin Dynabeads (Invitrogen) and washed twice with a low salt wash buffer (50mM Tris-HCl pH 7.5, 150mM NaCl) and twice with a high salt wash buffer (50mM Tris-HCl pH 7.5, 400mM NaCl) before being submitted to the Rockefeller University Proteomics Resource Center.

### RNA stability of hnRNPA1 bound transcripts

950,000 MCF10A cells were reverse transfected with a control siRNA or siRNA targeting hnRNPA1 with either a tRF^Tyr^_GUA_ mimetic or a scrambled mimetic using Lipofectamine RNAimax (Invitrogen). After 48 hours, half of the samples were treated with α-amanitin (10mg/ml, Sigma) and incubated for 8 hours while the other half were processed as the 0 hour time-point. RNA was extracted using TRIzol and RNA-seq libraries were prepared using QuantSeq 3’ mRNA-seq kit (Lexogen) following the manufacturer’s instructions.

For the converse experiment, siRNA was reverse transfected as above into 1,000,000 MCF10A cells using Lipofectamine RNAimax except a miRNA inhibitor (LNA) targeting tRF^Tyr^_GUA_ (GAUAGCUCAGUUGGUAGAGCGGAGGA)or non-targeting control (UCGUUAAUCGGCUAUAAUACGC) was directly transfected 24 hours later. After 48 hours, half of the samples were treated with α-amanitin (10mg/ml, Sigma) and 200µM H_2_O_2_ and incubated for 8 hours while the other half were processed as the 0 hour time-point. RNA was extracted using TRIzol and RNA-seq libraries were prepared using TruSeq RNA Library Prep (Illumina) following the manufacturer’s instructions.

### SSB HITS-CLIP with and without RNase

HITS-CLIP for endogenous SSB was done as described by (Licatalosi et al., 2008) with the modifications previously used for YBX1 small RNA CLIP (Goodarzi et al., 2015). MDA-MB-231 cells were UV-crosslinked at 400 mJ/cm^2^ before cell lysis. Samples with and without RNase treatment were immunoprecipitated with an anti-SSB antibody (MBL, RN074PW) for protein-RNA complexes. Polyphosphatase (Lucigen) was incubated with smRNA samples before ligation and PCR amplification with primers described by (Goodarzi et al., 2015). Constructed libraries were sequenced on the Illumina HiSeq2000 at the Rockefeller University Genomics Center.

The CTK package (Shah et al., 2017) was used to analyze the SSB HITS-CLIP data. First, reads were distributed to individual samples based on the integrated 4nt barcodes. The reads were then collapsed, trimmed, and aligned to the human genome (hg19) using bwa (0.7.17-r1188; -n 0.06 -q 20). The CTK toolkit was then used parse the alignments, remove PCR duplicates, and call mutations in reads. The results from biological replicates were then combined and the CIMS tool (CTK) was used to call peaks (FDR<0.1).

### Generation of CRISPRi-DIS3L2 knockdown cell lines

A non-targeting control sgRNA (sequence retrieved from hCRISPRi v2 (Horlbeck et al., 2016)) and 5 sgRNAs targeting *DIS3L2* (generated using the ChopChop tool (Labun et al., 2019)) were cloned into lentiGuide-Puro (Addgene #52963). The individual sgRNAs were subsequently transduced into MCF10a cells stably expressing pHR-SFFV-dCas9-BFP-KRAB (Addgene #46911) and the cells expressing the sgRNAs were enriched for using 1.2 ug/mL puromycin selection. The cells with the strongest knockdown of *DIS3L2* (>90%) by western blot were subsequently used for experiments, with the following guide sequences: sgRNA-NC1: GCGTACGACAATACGCGCGA, sgRNA-DIS3L2-A: CGCGGCGTTCTAGAGAGCGA.

### *C. elegans* culture conditions

Wild-type Bristol N2, and *disl-2 (syb1033)* mutant strains were cultured on nematode growth media (NGM) containing *E. coli* HB101 at 20°C. Synchronized populations of developmentally staged worms were obtained by standard methods. Post synchronization, worms were seeded on NGM plates and fed E. coli HB101 until they reached young adult stage. The *E. coli* HT115(DE3) strain was used for RNAi experiments.

To induce oxidative stress, approximately 3600 young adult worms were plated into 6-well plates and given a single dose of 200µM hydrogen peroxide in M9 buffer. Plates were rocked for 15 minutes prior to collection.

### *C. elegans* RNAi

Feeding RNAi bacterial clones carrying RNAi feeder plasmids of disl-2 were obtained from *C. elegans* RNAi v1.1 Feeding Library (Open Biosystems, Huntsville, AL, USA). Empty vector (pL4440) was used as a control. RNAi colonies for *disl-2* and *EV* control were grown overnight at 37°C with constant shaking in Luria broth with 100 μg/ml ampicillin (Sigma-Aldrich). The next day, these overnight cultures were seeded on RNAi plates containing 1mM isopropylthiogalactoside (IPTG, Fermentas) and 100 μg/ml ampicillin (Sigma-Aldrich) to induce dsRNA expression.

### *C. elegans Disl-2(syb1033)* PHX1033 mutant generation and genotyping

The allele harboring a mutation was generated using CRISPR method in N2 strain (Sunybiotech). sgRNA target sequence CCTCACTCAGACAGCTACAGGCG was used, and non-synonymous mutations were made in the sgRNA/PAM site. The wild-type *disl-2* sequence used was GTTGAAGCCGCAGGGC[<u>CTC]</u>ACTCAGACAGCTACAGG, and a *syb1033* mutant sequence of a CTC to TAG (GTTGAAGCCGCAGGGC[<u>TAG]</u>–CTCAGACAGCTACAGG). Mutants were confirmed by genotyping using PCR amplification with the following primers GATTTTACCCGACAGCGAT; TTACAGCCACCACATCTCCT. Products were sequence-verified and visualized on a gel following Nhe1-mediated restriction digestion.

## ACCESSION NUMBERS

The data for high-throughput sequencing experiments are deposited at GEO under the accession number GSE120385, with the following reviewer access token: udevwcychngxlsn

## AUTHOR CONTRIBUTIONS

S.F.T. conceived the project and supervised all research. S.F.T and D.H. wrote the manuscript. D.H., M.C.P., J.G., S.D., C.G., L.F., A.P., H.M., E.A.M., and H.G designed, performed, and analyzed the experiments.

## ACKNOWLEDGEMENTS

We are grateful to the members of our lab for their insightful comments on past versions of this manuscript. We thank Josh Mendell for generously providing the HEK293T DIS3L2 knockout cells. We thank Cori Bargmann and the Bargmann lab for generous help on *C. elegans* work as well as providing the RNAi clones used for this work. We are grateful to C. Zhao, C. Lai, and N. Nnatubeugo of the Rockefeller Genomics Resource Center for assistance with next-generation sequencing. D.H. and M.C.P. were supported by a Medical Scientist Training Program grant from the National Institute of General Medical Sciences of the National Institutes of Health under award number T32GM007739 to the Weill Cornell/Rockefeller/Sloan Kettering Tri-Institutional MD-PhD program. M.C.P was supported by a F30 Predoctoral Fellowship from the National Cancer Institute of the National Institutes of Health under award number 1F30CA247026-01. E.A.M was supported by NIH training grant T32 CA 9673-39. The research of S.F.T. was supported in part by a Faculty Scholar grant from the Howard Hughes Medical Institute and by the DOD Collaborative Scholars and Innovators Award (grant W81XWH-12-1-0301), Pershing Square Sohn award, Breast Cancer Research Foundation award, Reem-Kayden award, NIH grant 5R01CA215491-02, and the Black Family Metastasis Center. The content of this study is solely the responsibility of the authors and does not necessarily represent the official views of the National Institutes of Health.

---

**Figure S1. The fragmentation of tRNA^Tyr^_GUA_ in response to oxidative stress is not due to cell death.**

(A) The viability of MCF10A cells after H_2_O_2_ treatment (200μM) was tested by a trypan blue exclusion assay (n=6). Viability was tested one hour after exposure to oxidative stress. A two-tailed Mann-Whitney test was used to test for statistical significance between the treated and control cell lines for each time point.

(B) The log_2_ fold induction of tRFs for all isoacceptors in the overlapping set of tRNAs found in Fig 1c. Data represent mean ± s.e.m.

**Figure S2. Fragmentation of tRNA^Tyr^_GUA_ occurs with other sources of oxidative stress and additional cell lines.**

(A) Northern blot for tRF^Tyr^_GUA_ and tRF^Leu^_HAG_ in MCF10A cells at one hour post oxidative stress (200μM H_2_O_2_) exposure.

(B) Northern blot for tRF^Tyr^_GUA_ in C. elegans cells at 15 minutes post-oxidative stress (200μM H_2_O_2_) exposure.

(C) A northern blot depicting a time course experiment ranging from five minutes to 24 hours for HBEC30 cells in response to oxidative stress (200μM H_2_O_2_). As before, a single probe complementary to pre-tRNA^Tyr^_GUA_, mature tRNA^Tyr^_GUA_, and tRF^Tyr^_GUA_ was ^32^P-labeled and used for detection.

(D) A northern blot depicting two time points, one hour and 24 hours, after exposure to oxidative stress (200μM H_2_O_2_) in MCF10A cells. As before, a single probe complementary to pre-tRNA^Tyr^_GUA_, mature tRNA^Tyr^_GUA_, and tRF^Tyr^_GUA_ was while another probe complementary to either the mature tRNA^His^_GUG_ or mature tRNA^Glu^_YUC_ were both ^32^P-labeled and used for detection.

(E) Quantification of tRNA^Tyr^_GUA_, tRNA^His^_GUG_, and tRNA^Glu^_YUC_ by northern blot analysis from two independent experiments 24 hours (normalized to U6 levels) after exposure to oxidative stress (200μM H_2_O_2_) are shown (n=4). A one-tailed Mann-Whitney test was used to test for statistical significance between the treated and control cell lines for each time point.

(F) A northern blot at a short and longer time point after MCF10A cells were treated with a pharmacological agent used to induce oxidative stress (menadione).

(G) Quantification of tRNA^Tyr^_GUA_ by northern blot analysis from two independent experiments 24 hours (normalized to U6 levels) after exposure to oxidative stress (50μM menadione) are shown (n=4). A one-tailed Mann-Whitney test was used to test for statistical significance Data represent mean ± s.e.m. *p < 0.05

**Figure S3. Validation of YARS knockdown and tRNA^Tyr^_GUA_ overexpression**

(A) Total mRNA from MCF10A cells stably expressing a short hairpin targeting YARS was analyzed by quantitative RT-PCR. The two cell lines with the best knockdown were used for subsequent experiments. A one-tailed Mann-Whitney test (*p < 0.05) was used to test for statistical significance between the treated and control cell lines for each time point.

(B) Northern blot of MCF10A cells transiently transfected with a tRNA^Tyr^_GUA_ overexpression vector relative to an empty control vector. TRNA^Tyr^_GUA_ overexpression is shown at 24, 48, and 72 hours post transfection.

Data represent mean ± s.e.m.

**Figure S4. Proteomic analysis identifies a tRNA^Tyr^_GUA_-dependent gene-set.**

(A) A plot showing the correlation between protein abundance changes in the proteome upon either tRNA^Tyr^_GUA_ depletion or YARS depletion (shYARS-2) relative to control cells. A Pearson’s two-sided test was used to test determine the statistical significance of the correlation between tRNA^Tyr^_GUA_ and YARS depletion effects across the detected proteome.

(B) A schematic showing locations of all tyrosine residues in the coding sequence of three target genes, EPCAM, SCD, and USP3.

(C) Levels of mRNA expression for target genes in cells depleted of either tRNA^Tyr^_GUA_ or YARS as measured by qRT-PCR (n=4). A one-tailed Mann-Whitney test was used to test for statistical significance between the knockdown and control cell lines’ gene expression values.

(D) Total mRNA from MCF10A cells transiently transfected with two independent siRNA targeting EPCAM, SCD, or USP3 was analyzed by quantitative RT-PCR at the end of each growth assay. A one-tailed Mann-Whitney test was used to establish statistical significance (n=3 except for siSCD-2 has n=2).

Data represent mean ± s.e.m. *p < 0.05

**Figure S5. Proteomic and ribosomal profiling validation of tRNA^Tyr^_GUA_-depleted cells.**

(A) MRNA expression levels for target genes 24 hours after treatment with H_2_O_2_ (200μM) as measured by qRT-PCR (n=9). A one-tailed Mann-Whitney test was used to test for statistical significance between the treated and control cell lines’ gene expression values.

(B) Examples of the mapped position of the 5’-end of reads near the start (top) or stop (bottom) codons are shown, revealing the characteristic 3-nucleotide periodicity of ribosomal positioning observed as previously described by (Ingolia et al., 2009).

(C) Histogram of the read length distribution of ribosome protected fragments observed upon ribosome profiling sequence analysis.

(D) Codon usage scatter plots of the ribosomal A-site of shTyr vs shControl from replicate ribosomal profiling experiments. The five most affected codons are labeled with a codon for tyrosine, UAC, being the most affected (red arrow). Gray outline illustrates the 95% confidence interval around the regression line.

**Figure S6. Characterization of the tRF^Tyr^_GUA_ and its potential functional role.**

(A) Length of 5’ leader sequence of all tRF^Tyr^_GUA_ by loci that were identified by sequencing.

(B) Sequence of each tRF^Tyr^_GUA_ by loci that were identified by sequencing. Conserved tRF^Tyr^_GUA_ found in each loci are highlighted in gray.

(C) A northern blot depicting the tRF^Tyr^_GUA_ induction in MCF10A cells with two independent siRNA each against angiogenin (ANG) or TSEN2 relative to a control siRNA.

(D) QRT-PCR validation of the siRNA-mediated knockdown of ANG relative to a control siRNA.

(E) QRT-PCR validation of the siRNA-mediated knockdown of TSEN2 relative to a control siRNA.

(F) A northern blot depicting the tRF^Tyr^_GUA_ induction in MCF10A cells with two independent siRNA against CLP1 relative to a control siRNA.

(G) QRT-PCR validation of the siRNA-mediated knockdown of CLP1 relative to a control siRNA.

(H) Cell growth of MCF10A cells transiently transfected with a tRF^Tyr^_GUA_ mimetic relative to a scrambled tRF control (n=3). A one-tailed Mann-Whitney test was used to test for significance at day 3.

(I) Western blot of downstream tRNA^Tyr^_GUA_-dependent genes 24 hours after transfection of tRF^Tyr^_GUA_ mimetic or a scrambled tRF control.

**Figure S7. Validation of interactions between RBPs with endogenous smRNAs.**

(A) Autoradiogram of endogenous SSB (red arrow) after immunoprecipitation from UV-crosslinked cells. ^32^P labeled ribonucleoprotein complexes were treated with either low or high concentrations of RNase A digestion prior to separation on SDS-PAGE gel.

(B-C) Autoradiograms of immunoprecipitations from cross-linked cells for endogenous hnRNPA1 (A) and hnRNPA0 (B). ^32^P labeled ribonucleoprotein complexes with either no, low, or high concentrations of RNase A digestion before separating on SDS-PAGE gels.

(D) IGV plots from the SSB HITS-CLIP reveals protein binding to the 3’ trailer end of pre-tRNAs arising from distinct tRNA loci.

(E) Quantitative RT-PCR expression levels for various Pol III transcribed targets from MCF10A cells transfected with the tRF^Tyr^_GUA_ mimetic or a scrambled control.

(F) MRNA expression levels for disl2 after RNAi treatment, as measured by qRT-PCR (n=3). A one-tailed Mann-Whitney test was used to test for statistical significance between control and knockdown gene expression values.

(G) A northern blot for tRF^Tyr^_GUA_ in wild type *N2* and homozygous *disl-2 (syb1033)* mutant C. elegans strains after 15 minutes of oxidative stress (200μM H_2_O_2_). A single probe complementary to *C. elegans* pre-tRNA^Tyr^_GUA_, mature tRNA^Tyr^_GUA_, and tRF^Tyr^_GUA_ was ^32^P-labeled and used for detection.

(H) A northern blot for tRF^Tyr^_GUA_ in wild type *N2* empty vector control and RNAi-mediated *disl-2* knockdown *C. elegans* strains after 15 minutes of oxidative stress (200μM H_2_O_2_). As before, a single probe complementary to *C. elegans* pre-tRNA^Tyr^_GUA_, mature tRNA^Tyr^_GUA_, and tRF^Tyr^_GUA_ was ^32^P-labeled and used for detection.

(I) Western blot showing CRISPRi-mediated depletion of DIS3L2 in MCF10A and HEK293T cells.

(J) A northern blot for tRF^Tyr^_GUA_ in DIS3L2 knockout HEK293T cells after 1 hour and 24 hours oxidative stress. As before, a single probe complementary to pre-tRNA^Tyr^_GUA_, mature tRNA^Tyr^_GUA_, and tRF^Tyr^_GUA_ was^32^P-labeled and used for detection. Right, quantification of tRNA^Tyr^_GUA_ after 1 hour of oxidative stress (n=1, two replicates, normalized to U6 levels).

(K) A northern blot for tRF^Tyr^_GUA_ in CRISPRi-mediated DIS3L2 knockdown MCF10A cells after 1 hour of oxidative stress. As before, a single probe complementary to pre-tRNA^Tyr^_GUA_, mature tRNA^Tyr^_GUA_, and tRF^Tyr^_GUA_ was ^32^P-labeled and used for detection. Right, quantification of tRNA^Tyr^_GUA_ by northern blot analysis. n=7 across three independent experiments (normalized to U6 levels). A one-tailed Mann-Whitney test was used to test for statistical significance.

Data represent mean ± s.e.m. *p < 0.05, **p < 0.01, and ***p < 0.001

## REFERENCES

Ashburner, M., Ball, C.A., Blake, J.A., Botstein, D., Butler, H., Cherry, J.M., Davis, A.P., Dolinski, K., Dwight, S.S., Eppig, J.T., et al. (2000). Gene ontology: tool for the unification of biology. The Gene Ontology Consortium. Nat Genet 25, 25–29.

Biamonti, G., Bassi, M.T., Cartegni, L., Mechta, F., Buvoli, M., Cobianchi, F., and Riva, S. (1993). Human hnRNP protein A1 gene expression. Structural and functional characterization of the promoter. J Mol Biol 230, 77–89.

Boel, G., Letso, R., Neely, H., Price, W.N., Wong, K.H., Su, M., Luff, J., Valecha, M., Everett, J.K., Acton, T.B., et al. (2016). Codon influence on protein expression in E. coli correlates with mRNA levels. Nature 529, 358–363.

Cervantes Gracia, K., Llanas-Cornejo, D., and Husi, H. (2017). CVD and Oxidative Stress. J Clin Med 6.

Chan, C.T., Dyavaiah, M., DeMott, M.S., Taghizadeh, K., Dedon, P.C., and Begley, T.J. (2010). A quantitative systems approach reveals dynamic control of tRNA modifications during cellular stress. PLoS Genet 6, e1001247.

Chen, Q., Yan, M., Cao, Z., Li, X., Zhang, Y., Shi, J., Feng, G.H., Peng, H., Zhang, X., Zhang, Y., et al. (2016). Sperm tsRNAs contribute to intergenerational inheritance of an acquired metabolic disorder. Science 351, 397–400.

Chi, S.W., Zang, J.B., Mele, A., and Darnell, R.B. (2009). Argonaute HITS-CLIP decodes microRNA-mRNA interaction maps. Nature 460, 479–486.

Costa-Mattioli, M., Svitkin, Y., and Sonenberg, N. (2004). La autoantigen is necessary for optimal function of the poliovirus and hepatitis C virus internal ribosome entry site in vivo and in vitro. Mol Cell Biol 24, 6861–6870.

Couvillion, M.T., Sachidanandam, R., and Collins, K. (2010). A growth-essential Tetrahymena Piwi protein carries tRNA fragment cargo. Genes & development 24, 2742–2747.

Cozen, A.E., Quartley, E., Holmes, A.D., Hrabeta-Robinson, E., Phizicky, E.M., and Lowe, T.M. (2015). ARM-seq: AlkB-facilitated RNA methylation sequencing reveals a complex landscape of modified tRNA fragments. Nat Methods 12, 879–884.

Crick, F.H. (1966). Codon--anticodon pairing: the wobble hypothesis. J Mol Biol 19, 548–555.

Cross, C.E., Halliwell, B., Borish, E.T., Pryor, W.A., Ames, B.N., Saul, R.L., McCord, J.M., and Harman, D. (1987). Oxygen radicals and human disease. Ann Intern Med 107, 526–545.

Dahlgren, C., and Karlsson, A. (1999). Respiratory burst in human neutrophils. J Immunol Methods 232, 3–14.

Dittmar, K.A., Goodenbour, J.M., and Pan, T. (2006). Tissue-specific differences in human transfer RNA expression. PLoS Genet 2, e221.

dos Reis, M., Savva, R., and Wernisch, L. (2004). Solving the riddle of codon usage preferences: a test for translational selection. Nucleic Acids Res 32, 5036–5044.

Fu, H., Feng, J., Liu, Q., Sun, F., Tie, Y., Zhu, J., Xing, R., Sun, Z., and Zheng, X. (2009). Stress induces tRNA cleavage by angiogenin in mammalian cells. FEBS letters 583, 437–442.

Gehrke, C.W., Kuo, K.C., Waalkes, T.P., and Borek, E. (1979). Patterns of urinary excretion of modified nucleosides. Cancer Res 39, 1150–1153.

Gingold, H., Tehler, D., Christoffersen, N.R., Nielsen, M.M., Asmar, F., Kooistra, S.M., Christophersen, N.S., Christensen, L.L., Borre, M., Sorensen, K.D., et al. (2014). A dual program for translation regulation in cellular proliferation and differentiation. Cell 158, 1281–1292.

Glisovic, T., Ben-David, Y., Lang, M.A., and Raffalli-Mathieu, F. (2003). Interplay between hnRNP A1 and a cis-acting element in the 3’ UTR of CYP2A5 mRNA is central for high expression of the gene. FEBS letters 535, 147–152.

Gogakos, T., Brown, M., Garzia, A., Meyer, C., Hafner, M., and Tuschl, T. (2017). Characterizing Expression and Processing of Precursor and Mature Human tRNAs by Hydro-tRNAseq and PAR-CLIP. Cell Rep 20, 1463–1475.

Goncalves, K.A., Silberstein, L., Li, S., Severe, N., Hu, M.G., Yang, H., Scadden, D.T., and Hu, G.F. (2016). Angiogenin Promotes Hematopoietic Regeneration by Dichotomously Regulating Quiescence of Stem and Progenitor Cells. Cell 166, 894–906.

Goodarzi, H., Elemento, O., and Tavazoie, S. (2009). Revealing global regulatory perturbations across human cancers. Mol Cell 36, 900–911.

Goodarzi, H., Liu, X., Nguyen, H.C., Zhang, S., Fish, L., and Tavazoie, S.F. (2015). Endogenous tRNA-Derived Fragments Suppress Breast Cancer Progression via YBX1 Displacement. Cell 161, 790–802.

Goodarzi, H., Nguyen, H.C.B., Zhang, S., Dill, B.D., Molina, H., and Tavazoie, S.F. (2016). Modulated Expression of Specific tRNAs Drives Gene Expression and Cancer Progression. Cell 165, 1416–1427.

Gottlieb, E., and Steitz, J.A. (1989). Function of the mammalian La protein: evidence for its action in transcription termination by RNA polymerase III. EMBO J 8, 851–861.

Gustafsson, C., Govindarajan, S., and Minshull, J. (2004). Codon bias and heterologous protein expression. Trends Biotechnol 22, 346–353.

Guzzi, N., Ciesla, M., Ngoc, P.C.T., Lang, S., Arora, S., Dimitriou, M., Pimkova, K., Sommarin, M.N.E., Munita, R., Lubas, M., et al. (2018). Pseudouridylation of tRNA-Derived Fragments Steers Translational Control in Stem Cells. Cell 173, 1204–1216 e1226.

Hafner, M., Landthaler, M., Burger, L., Khorshid, M., Hausser, J., Berninger, P., Rothballer, A., Ascano, M., Jr., Jungkamp, A.C., Munschauer, M., et al. (2010). Transcriptome-wide identification of RNA-binding protein and microRNA target sites by PAR-CLIP. Cell 141, 129–141.

Hanada, T., Weitzer, S., Mair, B., Bernreuther, C., Wainger, B.J., Ichida, J., Hanada, R., Orthofer, M., Cronin, S.J., Komnenovic, V., et al. (2013). CLP1 links tRNA metabolism to progressive motor-neuron loss. Nature 495, 474–480.

Haussecker, D., Huang, Y., Lau, A., Parameswaran, P., Fire, A.Z., and Kay, M.A. (2010). Human tRNA-derived small RNAs in the global regulation of RNA silencing. RNA 16, 673–695.

Hoekema, A., Kastelein, R.A., Vasser, M., and de Boer, H.A. (1987). Codon replacement in the PGK1 gene of Saccharomyces cerevisiae: experimental approach to study the role of biased codon usage in gene expression. Mol Cell Biol 7, 2914–2924.

Honda, S., Loher, P., Shigematsu, M., Palazzo, J.P., Suzuki, R., Imoto, I., Rigoutsos, I., and Kirino, Y. (2015). Sex hormone-dependent tRNA halves enhance cell proliferation in breast and prostate cancers. Proc Natl Acad Sci U S A 112, E3816–3825.

Hopper, A.K., and Nostramo, R.T. (2019). tRNA Processing and Subcellular Trafficking Proteins Multitask in Pathways for Other RNAs. Front Genet 10, 96.

Horlbeck, M.A., Gilbert, L.A., Villalta, J.E., Adamson, B., Pak, R.A., Chen, Y., Fields, A.P., Park, C.Y., Corn, J.E., Kampmann, M., et al. (2016). Compact and highly active next-generation libraries for CRISPR-mediated gene repression and activation. Elife 5.

Huelga, S.C., Vu, A.Q., Arnold, J.D., Liang, T.Y., Liu, P.P., Yan, B.Y., Donohue, J.P., Shiue, L., Hoon, S., Brenner, S., et al. (2012). Integrative genome-wide analysis reveals cooperative regulation of alternative splicing by hnRNP proteins. Cell Rep 1, 167–178.

Hunter, R.W., Liu, Y., Manjunath, H., Acharya, A., Jones, B.T., Zhang, H., Chen, B., Ramalingam, H., Hammer, R.E., Xie, Y., et al. (2018). Loss of Dis3l2 partially phenocopies Perlman syndrome in mice and results in up-regulation of Igf2 in nephron progenitor cells. Genes & development 32, 903–908.

Ingolia, N.T., Ghaemmaghami, S., Newman, J.R., and Weissman, J.S. (2009). Genome-wide analysis in vivo of translation with nucleotide resolution using ribosome profiling. Science 324, 218–223.

Ishimura, R., Nagy, G., Dotu, I., Zhou, H., Yang, X.L., Schimmel, P., Senju, S., Nishimura, Y., Chuang, J.H., and Ackerman, S.L. (2014). RNA function. Ribosome stalling induced by mutation of a CNS-specific tRNA causes neurodegeneration. Science 345, 455–459.

Jean-Philippe, J., Paz, S., and Caputi, M. (2013). hnRNP A1: the Swiss army knife of gene expression. Int J Mol Sci 14, 18999–19024.

Karaca, E., Weitzer, S., Pehlivan, D., Shiraishi, H., Gogakos, T., Hanada, T., Jhangiani, S.N., Wiszniewski, W., Withers, M., Campbell, I.M., et al. (2014). Human CLP1 mutations alter tRNA biogenesis, affecting both peripheral and central nervous system function. Cell 157, 636–650.

Keam, S.P., Sobala, A., Ten Have, S., and Hutvagner, G. (2017). tRNA-Derived RNA Fragments Associate with Human Multisynthetase Complex (MSC) and Modulate Ribosomal Protein Translation. J Proteome Res 16, 413–420.

Kim, H.K., Fuchs, G., Wang, S., Wei, W., Zhang, Y., Park, H., Roy-Chaudhuri, B., Li, P., Xu, J., Chu, K., et al. (2017). A transfer-RNA-derived small RNA regulates ribosome biogenesis. Nature 552, 57–62.

Kuscu, C., Kumar, P., Kiran, M., Su, Z., Malik, A., and Dutta, A. (2018). tRNA fragments (tRFs) guide Ago to regulate gene expression post-transcriptionally in a Dicer independent manner. RNA.

Labun, K., Montague, T.G., Krause, M., Torres Cleuren, Y.N., Tjeldnes, H., and Valen, E. (2019). CHOPCHOP v3: expanding the CRISPR web toolbox beyond genome editing. Nucleic Acids Res 47, W171–W174.

Ladner, J.E., Jack, A., Robertus, J.D., Brown, R.S., Rhodes, D., Clark, B.F., and Klug, A. (1975). Structure of yeast phenylalanine transfer RNA at 2.5 A resolution. Proc Natl Acad Sci U S A 72, 4414–4418.

Lareau, L.F., Hite, D.H., Hogan, G.J., and Brown, P.O. (2014). Distinct stages of the translation elongation cycle revealed by sequencing ribosome-protected mRNA fragments. Elife 3, e01257.

Lee, S.R., and Collins, K. (2005). Starvation-induced cleavage of the tRNA anticodon loop in Tetrahymena thermophila. The Journal of biological chemistry 280, 42744–42749.

Lee, Y.S., Shibata, Y., Malhotra, A., and Dutta, A. (2009). A novel class of small RNAs: tRNA-derived RNA fragments (tRFs). Genes & development 23, 2639–2649.

Licatalosi, D.D., Mele, A., Fak, J.J., Ule, J., Kayikci, M., Chi, S.W., Clark, T.A., Schweitzer, A.C., Blume, J.E., Wang, X., et al. (2008). HITS-CLIP yields genome-wide insights into brain alternative RNA processing. Nature 456, 464–469.

Lin, M.T., and Beal, M.F. (2006). Mitochondrial dysfunction and oxidative stress in neurodegenerative diseases. Nature 443, 787–795.

Maraia, R.J., Kenan, D.J., and Keene, J.D. (1994). Eukaryotic transcription termination factor La mediates transcript release and facilitates reinitiation by RNA polymerase III. Mol Cell Biol 14, 2147–2158.

Maraia, R.J., Mattijssen, S., Cruz-Gallardo, I., and Conte, M.R. (2017). The La and related RNA-binding proteins (LARPs): structures, functions, and evolving perspectives. Wiley interdisciplinary reviews RNA 8.

Martindale, J.L., and Holbrook, N.J. (2002). Cellular response to oxidative stress: signaling for suicide and survival. J Cell Physiol 192, 1–15.

Mattijssen, S., Arimbasseri, A.G., Iben, J.R., Gaidamakov, S., Lee, J., Hafner, M., and Maraia, R.J. (2017). LARP4 mRNA codon-tRNA match contributes to LARP4 activity for ribosomal protein mRNA poly(A) tail length protection. Elife 6.

McGlincy, N.J., and Ingolia, N.T. (2017). Transcriptome-wide measurement of translation by ribosome profiling. Methods 126, 112–129.

Mili, S., and Steitz, J.A. (2004). Evidence for reassociation of RNA-binding proteins after cell lysis: implications for the interpretation of immunoprecipitation analyses. RNA 10, 1692–1694.

Munz, M., Baeuerle, P.A., and Gires, O. (2009). The emerging role of EpCAM in cancer and stem cell signaling. Cancer Res 69, 5627–5629.

Nicassio, F., Corrado, N., Vissers, J.H., Areces, L.B., Bergink, S., Marteijn, J.A., Geverts, B., Houtsmuller, A.B., Vermeulen, W., Di Fiore, P.P., et al. (2007). Human USP3 is a chromatin modifier required for S phase progression and genome stability. Curr Biol 17, 1972–1977.

Nissen, P., Hansen, J., Ban, N., Moore, P.B., and Steitz, T.A. (2000). The structural basis of ribosome activity in peptide bond synthesis. Science 289, 920–930.

Ogle, J.M., Murphy, F.V., Tarry, M.J., and Ramakrishnan, V. (2002). Selection of tRNA by the ribosome requires a transition from an open to a closed form. Cell 111, 721–732.

Parisien, M., Wang, X., and Pan, T. (2013). Diversity of human tRNA genes from the 1000-genomes project. RNA biology 10, 1853–1867.

Paton, C.M., and Ntambi, J.M. (2009). Biochemical and physiological function of stearoyl-CoA desaturase. Am J Physiol Endocrinol Metab 297, E28–37.

Paushkin, S.V., Patel, M., Furia, B.S., Peltz, S.W., and Trotta, C.R. (2004). Identification of a human endonuclease complex reveals a link between tRNA splicing and pre-mRNA 3’ end formation. Cell 117, 311–321.

Pavon-Eternod, M., Gomes, S., Geslain, R., Dai, Q., Rosner, M.R., and Pan, T. (2009). tRNA over-expression in breast cancer and functional consequences. Nucleic Acids Res 37, 7268–7280.

Pavon-Eternod, M., Gomes, S., Rosner, M.R., and Pan, T. (2013). Overexpression of initiator methionine tRNA leads to global reprogramming of tRNA expression and increased proliferation in human epithelial cells. RNA 19, 461–466.

Pershing, N.L., Lampson, B.L., Belsky, J.A., Kaltenbrun, E., MacAlpine, D.M., and Counter, C.M. (2015). Rare codons capacitate Kras-driven de novo tumorigenesis. J Clin Invest 125, 222–233.

Piskounova, E., Agathocleous, M., Murphy, M.M., Hu, Z., Huddlestun, S.E., Zhao, Z., Leitch, A.M., Johnson, T.M., DeBerardinis, R.J., and Morrison, S.J. (2015). Oxidative stress inhibits distant metastasis by human melanoma cells. Nature 527, 186–191.

Presnyak, V., Alhusaini, N., Chen, Y.H., Martin, S., Morris, N., Kline, N., Olson, S., Weinberg, D., Baker, K.E., Graveley, B.R., et al. (2015). Codon optimality is a major determinant of mRNA stability. Cell 160, 1111–1124.

Quigley, G.J., and Rich, A. (1976). Structural domains of transfer RNA molecules. Science 194, 796–806.

Radi, R. (2018). Oxygen radicals, nitric oxide, and peroxynitrite: Redox pathways in molecular medicine. Proc Natl Acad Sci U S A 115, 5839–5848.

Roy, R., Huang, Y., Seckl, M.J., and Pardo, O.E. (2017). Emerging roles of hnRNPA1 in modulating malignant transformation. Wiley interdisciplinary reviews RNA 8.

Saikia, M., Krokowski, D., Guan, B.J., Ivanov, P., Parisien, M., Hu, G.F., Anderson, P., Pan, T., and Hatzoglou, M. (2012). Genome-wide identification and quantitative analysis of cleaved tRNA fragments induced by cellular stress. The Journal of biological chemistry 287, 42708–42725.

Schaffer, A.E., Eggens, V.R., Caglayan, A.O., Reuter, M.S., Scott, E., Coufal, N.G., Silhavy, J.L., Xue, Y., Kayserili, H., Yasuno, K., et al. (2014). CLP1 founder mutation links tRNA splicing and maturation to cerebellar development and neurodegeneration. Cell 157, 651–663.

Schimmel, P. (2018). The emerging complexity of the tRNA world: mammalian tRNAs beyond protein synthesis. Nat Rev Mol Cell Biol 19, 45–58.

Shah, A., Qian, Y., Weyn-Vanhentenryck, S.M., and Zhang, C. (2017). CLIP Tool Kit (CTK): a flexible and robust pipeline to analyze CLIP sequencing data. Bioinformatics 33, 566–567.

Sharma, U., Conine, C.C., Shea, J.M., Boskovic, A., Derr, A.G., Bing, X.Y., Belleannee, C., Kucukural, A., Serra, R.W., Sun, F., et al. (2016). Biogenesis and function of tRNA fragments during sperm maturation and fertilization in mammals. Science 351, 391–396.

The Gene Ontology, C. (2017). Expansion of the Gene Ontology knowledgebase and resources. Nucleic Acids Res 45, D331–D338.

Thompson, D.M., Lu, C., Green, P.J., and Parker, R. (2008). tRNA cleavage is a conserved response to oxidative stress in eukaryotes. RNA 14, 2095–2103.

Thompson, D.M., and Parker, R. (2009a). The RNase Rny1p cleaves tRNAs and promotes cell death during oxidative stress in Saccharomyces cerevisiae. J Cell Biol 185, 43–50.

Thompson, D.M., and Parker, R. (2009b). Stressing out over tRNA cleavage. Cell 138, 215–219.

Torrent, M., Chalancon, G., de Groot, N.S., Wuster, A., and Madan Babu, M. (2018). Cells alter their tRNA abundance to selectively regulate protein synthesis during stress conditions. Sci Signal 11.

Truitt, M.L., and Ruggero, D. (2016). New frontiers in translational control of the cancer genome. Nat Rev Cancer 16, 288–304.

Ule, J., Jensen, K.B., Ruggiu, M., Mele, A., Ule, A., and Darnell, R.B. (2003). CLIP identifies Nova-regulated RNA networks in the brain. Science 302, 1212–1215.

Wang, H., Han, L., Zhao, G., Shen, H., Wang, P., Sun, Z., Xu, C., Su, Y., Li, G., Tong, T., et al. (2016). hnRNP A1 antagonizes cellular senescence and senescence-associated secretory phenotype via regulation of SIRT1 mRNA stability. Aging Cell 15, 1063–1073.

Weitzer, S., and Martinez, J. (2007). The human RNA kinase hClp1 is active on 3’ transfer RNA exons and short interfering RNAs. Nature 447, 222–226.

Wohlgemuth, S.E., Gorochowski, T.E., and Roubos, J.A. (2013). Translational sensitivity of the Escherichia coli genome to fluctuating tRNA availability. Nucleic Acids Res 41, 8021–8033.

Wolin, S.L., and Cedervall, T. (2002). The La protein. Annu Rev Biochem 71, 375–403.

Wu, C.C., Zinshteyn, B., Wehner, K.A., and Green, R. (2019). High-Resolution Ribosome Profiling Defines Discrete Ribosome Elongation States and Translational Regulation during Cellular Stress. Mol Cell 73, 959–970 e955.

Xiao, Z., Zou, Q., Liu, Y., and Yang, X. (2016). Genome-wide assessment of differential translations with ribosome profiling data. Nat Commun 7, 11194.

Yamasaki, S., Ivanov, P., Hu, G.F., and Anderson, P. (2009). Angiogenin cleaves tRNA and promotes stress-induced translational repression. J Cell Biol 185, 35–42.

Yoo, C.J., and Wolin, S.L. (1997). The yeast La protein is required for the 3’ endonucleolytic cleavage that matures tRNA precursors. Cell 89, 393–402.

Zheng, G., Qin, Y., Clark, W.C., Dai, Q., Yi, C., He, C., Lambowitz, A.M., and Pan, T. (2015). Efficient and quantitative high-throughput tRNA sequencing. Nat Methods 12, 835–837.

Zhong, J., Xiao, C., Gu, W., Du, G., Sun, X., He, Q.Y., and Zhang, G. (2015). Transfer RNAs Mediate the Rapid Adaptation of Escherichia coli to Oxidative Stress. PLoS Genet 11, e1005302.

