## Supplemental Figures 1-7 for "A stress-induced Tyrosine tRNA depletion response mediates codon-based translational repression and growth suppression"

A

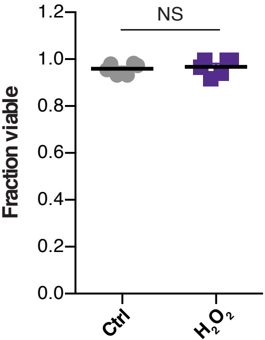


B

| **tRNA** | **$\text{Log}_2$ Fold change** |
|---|---|
| $\text{Tyr}^{\text{GUA}}$ | 1.16 |
| $\text{Leu}^{\text{CAA}}$ | 1.53 |
| $\text{Leu}^{\text{TAA}}$ | 2.19 |
| $\text{Leu}^{\text{CAG}}$ | 0.54 |
| $\text{Leu}^{\text{AAG}}$ | 0.75 |
| $\text{Leu}^{\text{TAG}}$ | 0.40 |
| $\text{Ile}^{\text{TAT}}$ | 1.56 |
| $\text{Ile}^{\text{AAT}}$ | 0.13 |
| $\text{Thr}^{\text{TGT}}$ | 0.38 |
| $\text{Thr}^{\text{AGT}}$ | 0.37 |

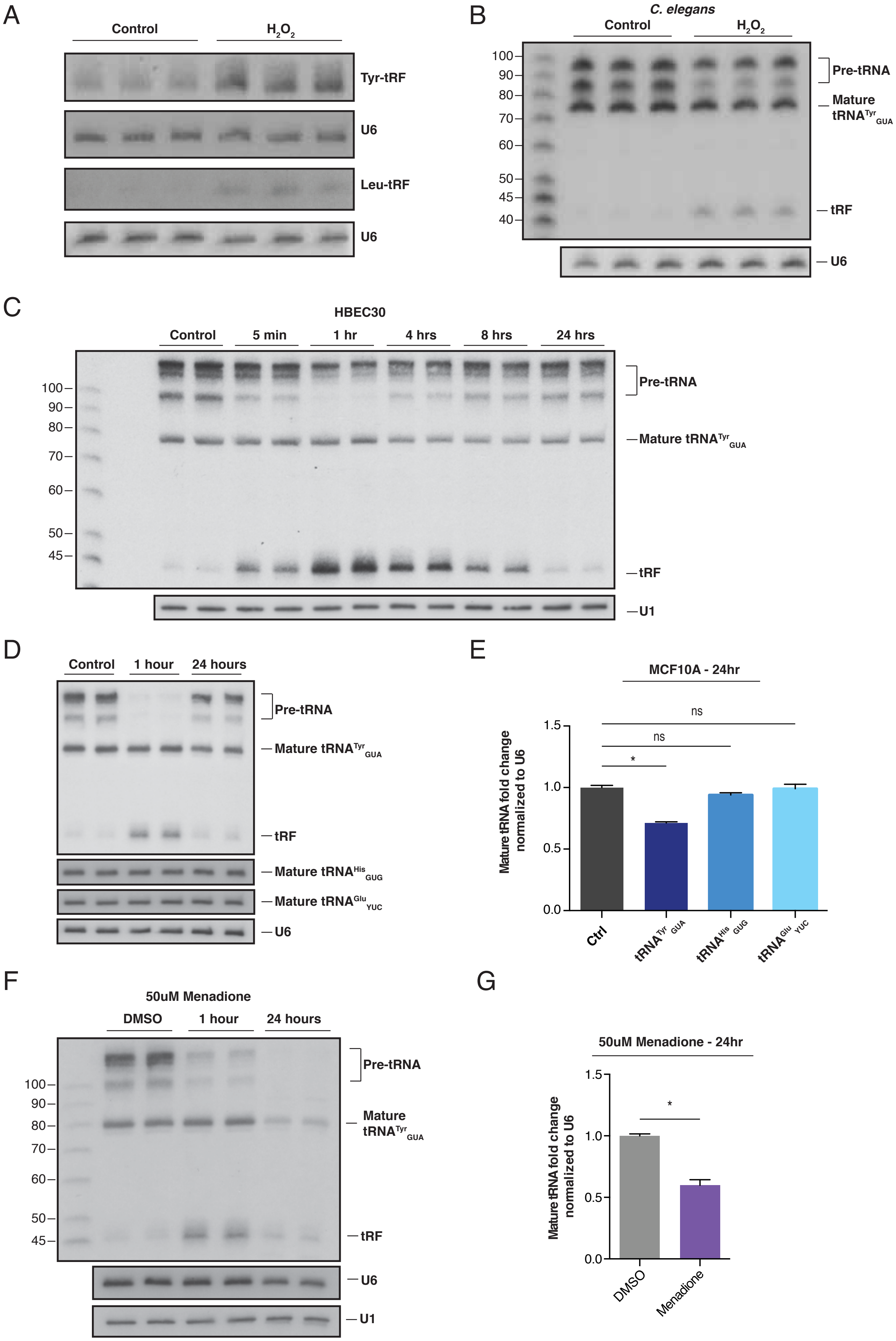
A
Control
H2O2
Tyr-tRF
U6
Leu-tRF
U6
B
C. elegans
Control
H2O2
100
90
80
70
60
50
45
40
Pre-tRNA
Mature tRNATyrGUA
tRF
U6
C
HBEC30
Control
5 min
1 hr
4 hrs
8 hrs
24 hrs
100
90
80
70
60
50
45
Pre-tRNA
Mature tRNATyrGUA
tRF
U1
D
Control
1 hour
24 hours
Pre-tRNA
Mature tRNATyrGUA
tRF
Mature tRNAHisGUG
Mature tRNAGluYUC
U6
E
MCF10A - 24hr
ns
ns
*
Mature tRNA fold change normalized to U6
1.5
1.0
0.5
0.0
Ctrl
tRNATyr GUA
tRNAHis GUG
tRNAGlu YUC
F
50uM Menadione
DMSO
1 hour
24 hours
100
90
80
70
60
50
45
Pre-tRNA
Mature tRNATyrGUA
tRF
U6
U1
G
50uM Menadione - 24hr
*
Mature tRNA fold change normalized to U6
1.5
1.0
0.5
0.0
DMSO
Menadione

A

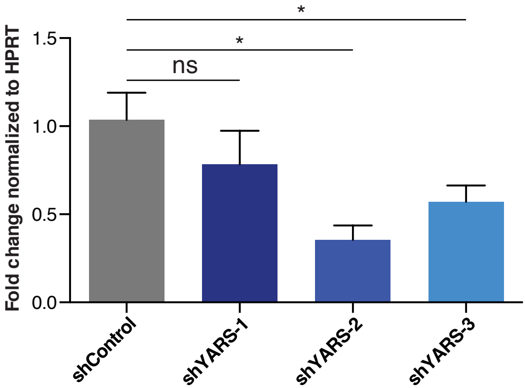
Fold change normalized to HPRT
1.5
1.0
0.5
0.0
ns
*
*
shControl
shYARS-1
shYARS-2
shYARS-3

B

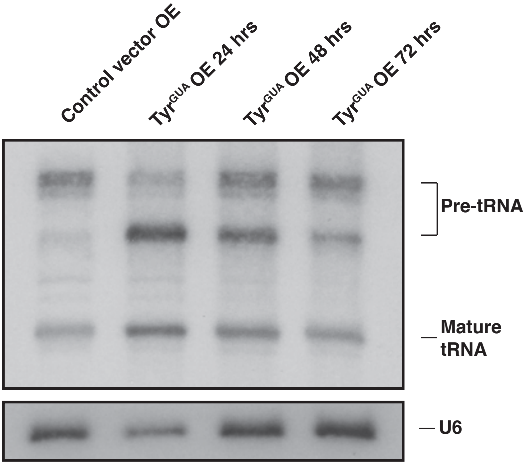
Control vector OE
Tyr GUA OE 24 hrs
Tyr GUA OE 48 hrs
Tyr GUA OE 72 hrs
Pre-tRNA
Mature tRNA
U6

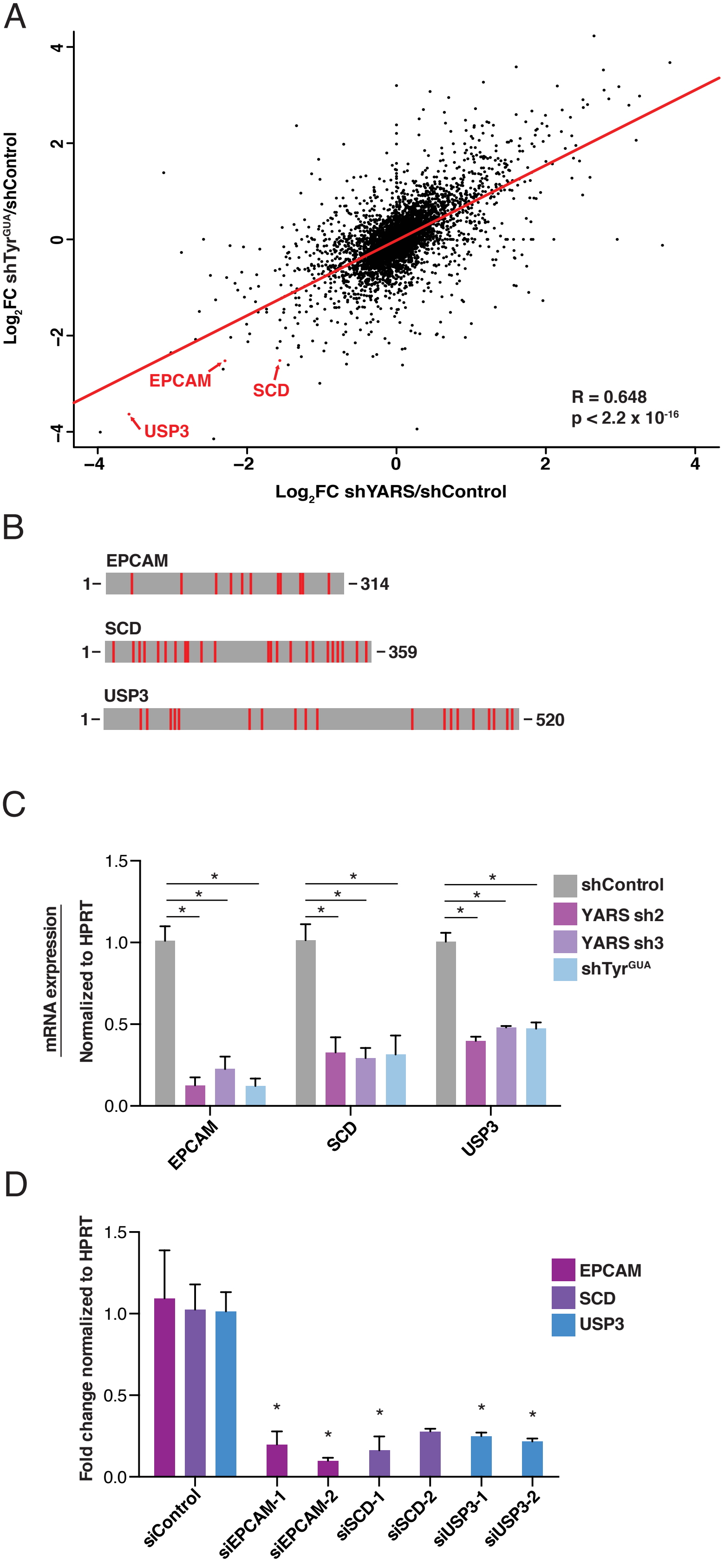
A
Log2FC shTyrGUA/shControl
Log2FC shYARS/shControl
EPCAM
SCD
USP3
R = 0.648
p < 2.2 x 10-16
B
EPCAM
1– –314
SCD
1– –359
USP3
1– –520
C
mRNA exrpression
Normalized to HPRT
shControl
YARS sh2
YARS sh3
shTyrGUA
EPCAM
SCD
USP3
D
Fold change normalized to HPRT
EPCAM
SCD
USP3
siControl
siEPCAM-1
siEPCAM-2
siSCD-1
siSCD-2
siUSP3-1
siUSP3-2

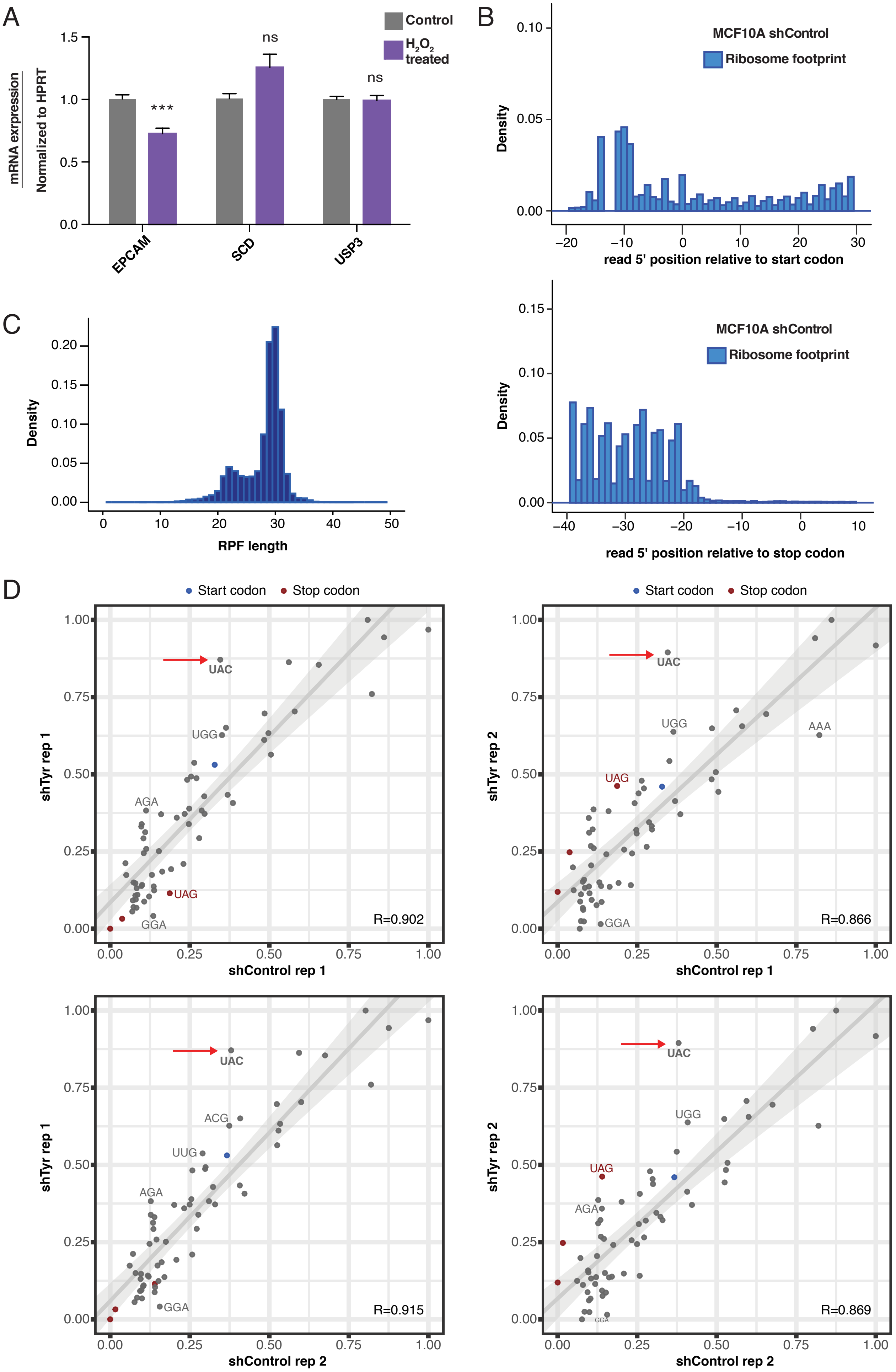

A
Control
$H_2O_2$ treated
mRNA exrpression
Normalized to HPRT
***
ns
ns
EPCAM
SCD
USP3
B
MCF10A shControl
Ribosome footprint
Density
read 5' position relative to start codon
MCF10A shControl
Ribosome footprint
Density
read 5' position relative to stop codon
C
Density
RPF length
D
Start codon
Stop codon
shTyr rep 1
shControl rep 1
UAC
UGG
AGA
UAG
GGA
R=0.902
shTyr rep 2
shControl rep 1
UAC
UGG
AAA
UAG
GGA
R=0.866
shTyr rep 1
shControl rep 2
UAC
ACG
UUG
AGA
GGA
R=0.915
shTyr rep 2
shControl rep 2
UAC
UGG
UAG
AGA
R=0.869

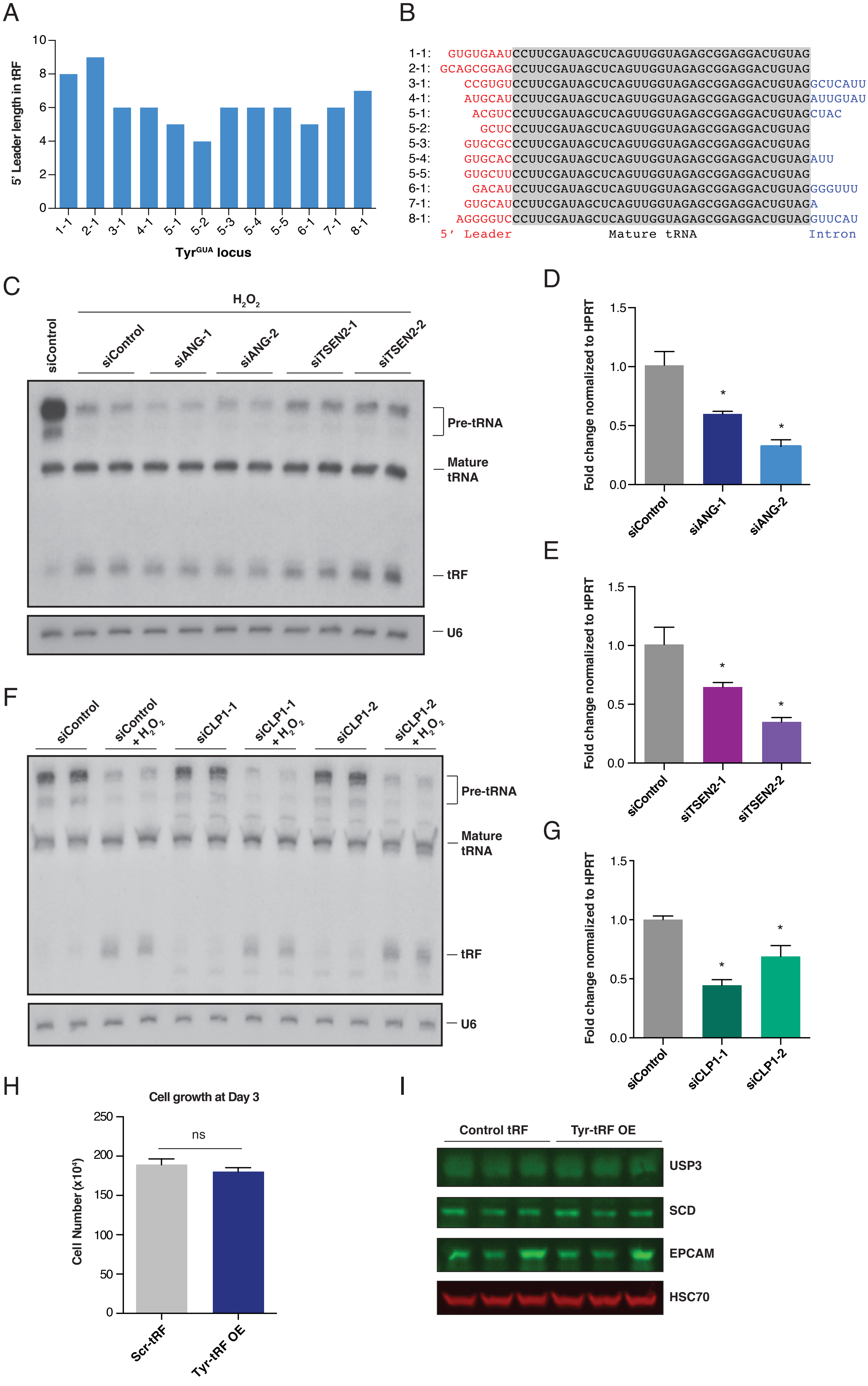

A
5' Leader length in tRF
Tyr^GUA locus
1-1 2-1 3-1 4-1 5-1 5-2 5-3 5-4 5-5 6-1 7-1 8-1
B
1-1: GUGUGAAUCCUUCGAUAGCUCAGUUGGUAGAGCGGAGGACUGUAG
2-1: GCAGCGGAGCCUUCGAUAGCUCAGUUGGUAGAGCGGAGGACUGUAG
3-1: CCGUGUCCUUCGAUAGCUCAGUUGGUAGAGCGGAGGACUGUAGGCUCAUU
4-1: AUGCAUCCUUCGAUAGCUCAGUUGGUAGAGCGGAGGACUGUAGAUUGUAU
5-1: ACGUCCCUUCGAUAGCUCAGUUGGUAGAGCGGAGGACUGUAGCUAC
5-2: GCUCCCUUCGAUAGCUCAGUUGGUAGAGCGGAGGACUGUAG
5-3: GUGCGCCCUUCGAUAGCUCAGUUGGUAGAGCGGAGGACUGUAG
5-4: GUGCACCCUUCGAUAGCUCAGUUGGUAGAGCGGAGGACUGUAGAUU
5-5: GUGCUUCCUUCGAUAGCUCAGUUGGUAGAGCGGAGGACUGUAG
6-1: GACAUCCUUCGAUAGCUCAGUUGGUAGAGCGGAGGACUGUAGGGGUUU
7-1: GUGCAUCCUUCGAUAGCUCAGUUGGUAGAGCGGAGGACUGUAGA
8-1: AGGGGUCCCUUCGAUAGCUCAGUUGGUAGAGCGGAGGACUGUAGGUUCAU
5' Leader
Mature tRNA
Intron
C
H2O2
siControl siControl siANG-1 siANG-2 siTSEN2-1 siTSEN2-2
Pre-tRNA
Mature tRNA
tRF
U6
D
Fold change normalized to HPRT
siControl siANG-1 siANG-2
E
Fold change normalized to HPRT
siControl siTSEN2-1 siTSEN2-2
F
siControl siControl + H2O2 siCLP1-1 siCLP1-1 + H2O2 siCLP1-2 siCLP1-2 + H2O2
Pre-tRNA
Mature tRNA
tRF
U6
G
Fold change normalized to HPRT
siControl siCLP1-1 siCLP1-2
H
Cell growth at Day 3
Cell Number (x10^4)
ns
Scr-tRF Tyr-tRF OE
I
Control tRF Tyr-tRF OE
USP3
SCD
EPCAM
HSC70

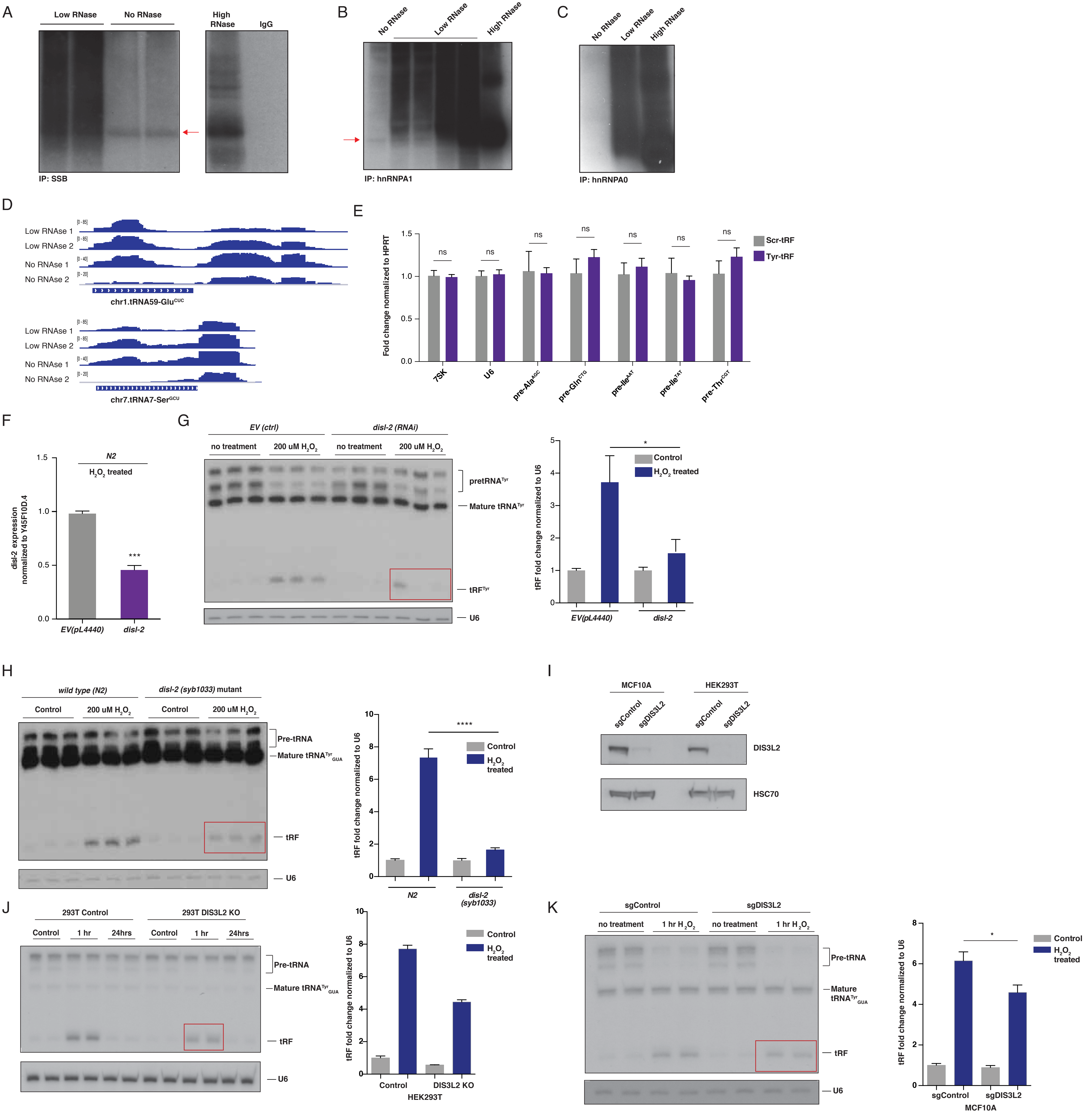
A
Low RNase
No RNase
High RNase
IgG
IP: SSB
B
No RNase
Low RNase
High RNase
IP: hnRNPA1
C
No RNase
Low RNase
High RNase
IP: hnRNPA0
D
Low RNAse 1
Low RNAse 2
No RNAse 1
No RNAse 2
[0 - 85]
[0 - 40]
[0 - 20]
chr1.tRNA59-Glu^CUC
chr7.tRNA7-Ser^GCU
E
Fold change normalized to HPRT
1.5
1.0
0.5
0.0
ns
Scr-tRF
Tyr-tRF
7SK
U6
pre-Ala^AGC
pre-Gln^CTG
pre-Ile^AAT
pre-Ile^TAT
pre-Thr^CGT
F
disl-2 expression normalized to Y45F10D.4
N2
H2O2 treated
***
EV(pL4440)
disl-2
G
EV (ctrl)
disl-2 (RNAi)
no treatment
200 uM H2O2
pretRNA^Tyr
Mature tRNA^Tyr
tRF^Tyr
U6
tRF fold change normalized to U6
*
Control
H2O2 treated
EV(pL4440)
disl-2
H
wild type (N2)
disl-2 (syb1033) mutant
Control
200 uM H2O2
Pre-tRNA
Mature tRNA^Tyr_GUA
tRF
U6
tRF fold change normalized to U6
****
Control
H2O2 treated
N2
disl-2 (syb1033)
I
MCF10A
HEK293T
sgControl
sgDIS3L2
DIS3L2
HSC70
J
293T Control
293T DIS3L2 KO
Control
1 hr
24hrs
Pre-tRNA
Mature tRNA^Tyr_GUA
tRF
U6
tRF fold change normalized to U6
Control
H2O2 treated
Control
DIS3L2 KO
HEK293T
K
sgControl
sgDIS3L2
no treatment
1 hr H2O2
Pre-tRNA
Mature tRNA^Tyr_GUA
tRF
U6
tRF fold change normalized to U6
*
Control
H2O2 treated
sgControl
sgDIS3L2
MCF10A
